# Multifunctional fibers enable modulation of cortical and deep brain activity during cognitive behavior in macaques

**DOI:** 10.1101/2022.10.09.511302

**Authors:** Indie C Garwood, Alex J Major, Marc-Joseph Antonini, Josefina Correa, Youngbin Lee, Atharva Sahasrabudhe, Meredith K Mahnke, Earl K Miller, Emery N Brown, Polina Anikeeva

## Abstract

Recording and modulating neural activity *in vivo* enables investigations of neural circuits during behavior. However, there is a dearth of tools for simultaneous recording and localized receptor modulation in large animal models. We address this limitation by translating multifunctional fiber-based neurotechnology previously only available for rodent studies to enable cortical and subcortical neural modulation in macaques. We record single unit and local field potential activity before, during, and after intracranial GABA infusions in the premotor cortex and putamen. We apply state-space models to characterize changes in neural activity and investigate how neural activity evoked by a working memory task varies in the presence of local inhibition. The recordings provide detailed insight into the electrophysiological effect of neurotransmitter receptor modulation in both cortical and subcortical structures in an awake, behaving macaque. Our results demonstrate a first-time translation of multifunctional fibers for causal studies in behaving non-human primates.

## 1 Main

Studying the relationship between neural activity and behavior is a cornerstone of systems and cognitive neuroscience research. Neural encoding of behavioral information occurs across multiple timescales and multiple modalities, including single neuron activity [1, 2], coordinated activity from populations of neurons [3], local field potential oscillations [4], and network communication across distant brain areas [5]. State-space and generalized linear models provide a powerful, flexible approach to decoding behavioral information from time-varying neurophysiology measurements [1, 6–9]. Together, observational studies and decoding advancements have enabled breakthroughs in brain machine interfaces for controlling prosthetic limbs and other assistive devices. However, it is challenging to establish causal rather than correlative associations between brain activity and behavior from a purely observational approach [10]. Establishing causal relationships is a critical step in understanding brain function and developing effective treatments for neurological and mental illnesses [11].

Recording the effect of neurophysiology perturbations can provide deeper insight into how neural activity encodes information [12]. While cell-type specific modulation of genetically engineered receptors is a powerful neuroscience research tool [13], genetic alterations are challenging in non-human primates (NHPs) [14], a critical translational model of higher cognitive function. Intracranial drug delivery enables modulation of specific receptors, and simultaneous electrophysiology measurements can establish causal links between receptor function and neural encoding [15–18]. Iontophoretic probes enable near simultaneous electrophysiology and drug delivery via application of electrical fields [17–21]. However, such devices are typically only used for delivery of polar molecules, and the applied field causes an electrical artifact in recordings [20]. Pressure injection with precise micropumps enables delivery of nanoliter volumes of a broader range of neuromodulator compounds and, at low flow rates, causes minimal recording artifacts [22, 23]. Consequently, several devices have been engineered to combine microfluidics with recording electrodes for NHPs [22–24]. These technologies, developed both in academic and commercial settings, employ rigid capillaries of steel or glass with diameters of 200–420 µm.

Multifunctional neurotechnology with smaller dimensions and softer materials (e.g., polymers and hydrogels) has the potential to reduce tissue damage and foreign body responses, enabling longer term experiments [25–29]. While the last decade of research into flexible bioelectronic interfaces has yielded an array of multifunctional tools for rodent studies [30, 31], few technologies developed for rodents have been translated to NHPs [32, 33]. Multifunctional fibers fabricated with the thermal drawing process provide an alternate paradigm for electrofluidic neurotechnology [34–37]. Thermally drawn fibers exhibit low bending stiffness due to miniaturization and the incorporation of flexible materials, which may reduce tissue damage caused by micromotion of the brain with respect to the implant and enable stable recording of single neuron activity [34–37]. The thermal drawing process yields meters of fiber in a single fabrication step, which are then sectioned into smaller segments as appropriate for the application. Thus, the thermal drawing process inherently enables scaling of flexible multifunctional probes from mouse to NHP applications.

Here, we present multifunctional fiber-based neural interfaces designed for NHP studies. The intracranial portion of the devices, the fiber, is 187.1± 2.5 µm in diameter and comprises four tungsten microelectrodes and two microfluidic channels integrated into a poly(etherimide) (PEI) cladding. We demonstrate the ability to locally modulate neural activity by delivering microinfusions of gamma-aminobutyric acid (GABA) to the premotor cortex of a rhesus macaque. We simultaneously record single unit and local field potential activity and demonstrate that GABA significantly alters neural activity. Because the premotor cortex is known to be involved in multiple cognitive processes, including working memory [38, 39], we leverage this experimental setup to explore how single unit and local field potential working memory encoding varies before and after local inhibition. Finally, we demonstrate that the probes are capable of recording and modulating task related activity in the putamen, a deep brain structure known to be involved in reward-mediated learning and working memory [40, 41]. The back-ends of the devices are designed for plug-and-play integration with existing experimental setups in labs using off-the-shelf NHP neurotechnologies. Consequently, we envision that these fiber-based tools will empower NHP neuroscience, enabling causal studies of the role of receptors in circuit function and high-level cognitive behaviors.

## 2 Results

### 2.1 Multifunctional fibers for drug delivery and electrophysiology

Although multifunctional fibers have been extensively applied for recording and modulation of neural activity in freely behaving rodents, the potential of this platform for multimodal neural interrogation in behaving NHPs remained to be harnessed. Here, we applied the thermal drawing process to create fibers capable of simultaneous drug delivery and electrophysiology recording in NHPs (Figure 1a). During thermal drawing, a macroscale preform is lowered into an oven and heated above its glass transition temperature, which causes the polymer to flow. The heated polymer is pulled downwards by a capstan, resulting in fiber with equivalent cross-sectional geometry but reduced dimensions (Figure 1b). We utilized PEI, a high performance thermoplastic, as the main structural component of our fiber. To incorporate tungsten electrodes into the fiber, we applied convergence drawing. The ends of four spools of tungsten microwires (diameter = 25 µm) were inserted into channels machined within the preform. As the fiber was drawn, the channels constricted around the wire, pulling the wires into the fiber. The diameter of the fiber was tuned by adjusting the relative rate that the preform was fed into the oven versus the rate the fiber was pulled out of the oven, producing 8 meters of fiber with diameter tuned between 150–200 µm.

**Figure 1:**
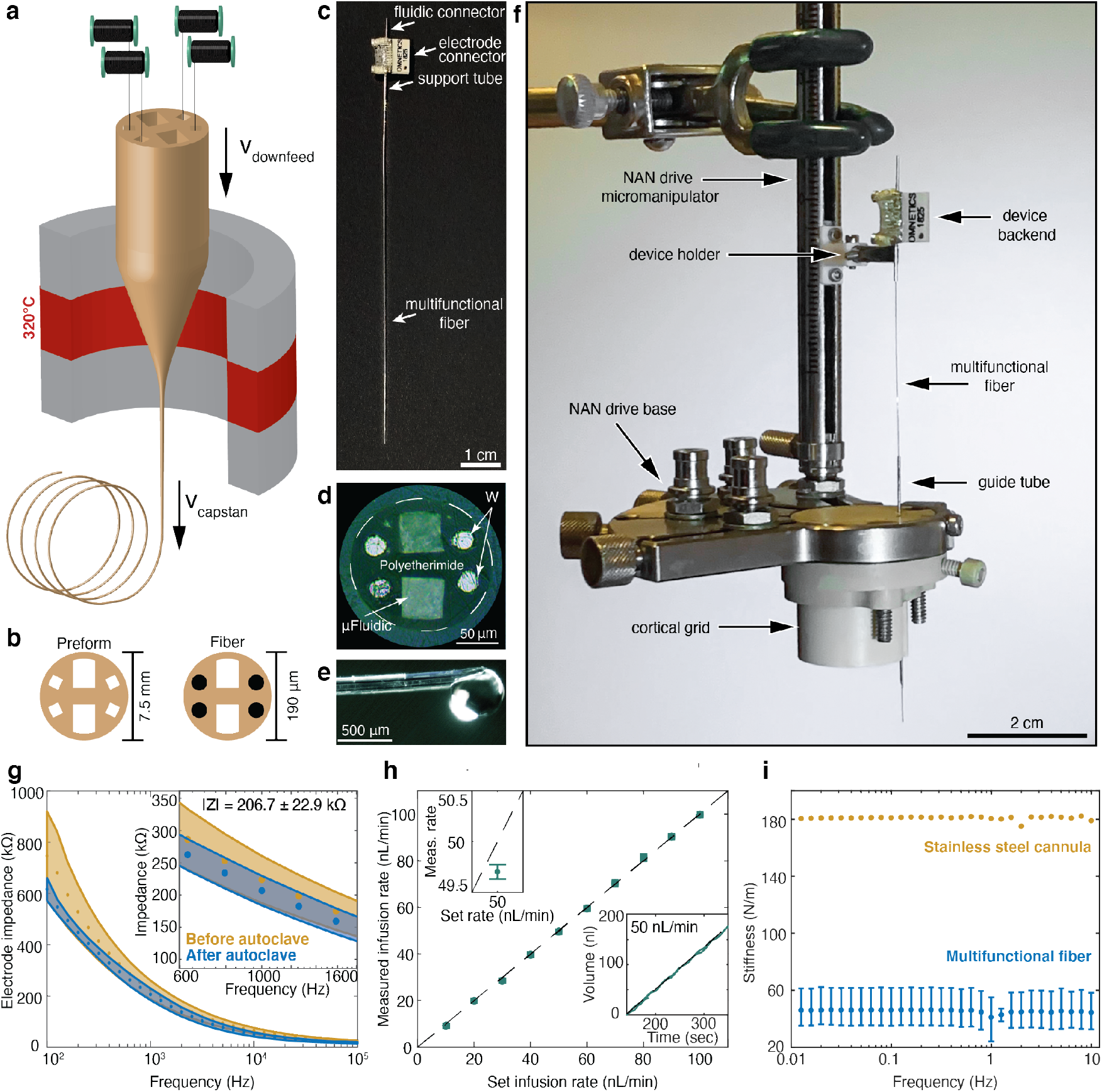
Fabrication of multifunctional fiber-based probes. **a**, The fiber preform was placed in a thermal drawing oven, where it was heated to 320°C. A capstan pulled downward on the preform at velocity *v*_capstan_ while the preform was lowered further into the oven at *v*_downfeed_. The cross-sectional area of the resulting fiber, *A*_fiber_ = (*V*_capstan_*/V*_downfeed_) *× A*_preform_. Tungsten (W) microwires were incorporated into the fiber via convergence (Methods). **b**, The resulting fiber (*d*_fiber_ = 187.1± 2.5 µm) has the same cross-sectional geometry as the preform (*d*_preform_ = 7.5 mm). **e**, Each device has an electrode connector for electrical interfacing, a stainless steel fluidic connector (ID = 304 µm, OD = 457 µm), a stainless steel support tube (ID = 432 µm, OD = 635 µm), and fiber (*d*_fiber_ = 187.1± 2.5 µm). We used fiber lengths of 7.0 ± 0.3 cm in this study, but lengths from 0–2 m are feasible. **d**, Cross section of fiber surrounded by embedding epoxy. The outside diameter of the fiber is indicated with the dashed white line. **e**, lateral view of a fiber tip. **f**, The devices were assembled in a cortical grid with a commercial microdrive. The cortical grid assembly includes a guide tube for dura penetration **g**, Impedance spectroscopy for the electrodes in the resulting fiber shows that the electrodes have a characteristic 1/f impedance curve over 10^2^ to 10^5^ Hz. The inset shows impedance between 600–1600 Hz. The mean impedances ± standard error before and after autoclave sterilization are shown in blue and yellow, respectively. Impedance at 1000 kHz = 223.9± 36.6 kΩ before autoclave sterilization and 206.7 ± 22.9 kΩ after. **h**, Characterization of the fluidic properties of the probe microfluidics demonstrates that the probes are capable of accurately injecting at rates of 10–100 nL/min. Each point shows the measured infusion rate and 95% confidence intervals; the top left inset shows the infusion rate error at 50 nL/min. The bottom right inset shows the infusion profile at steady state—volume flows at a near constant rate, and the mean absolute error (MAE) between the set volume and measured volume is 1.77 nL. **i**, Dynamic material analysis demonstrates that the fibers (*n* = 3) are less stiff than a stainless steel capillary (ID = 51 µm, OD = 203 µm OD).

Our devices were fabricated from 7.0 ± 0.3 cm sections of fiber with 187.1 ± 2.5 µm diameter (Figure 1c,d). The resulting fibers have four 25 µm tungsten microelectrodes and two (49.9 ± 0.7) × (62.4 ± 0.8) µm^2^ microfluidic channels (Figure 1d,e). The fiber length, which is approximately 25 times longer than those used for devices applied in rodent brains (Extended Data Figure 1a, [34]), enabled the devices to integrate with existing NHP microdrive setups (Figure 1f). The multifunctional fibers were outfitted with modular backends that employ commercial electrode connectors (Methods) and 304 µm inner diameter (ID), 457 µm outer diameter (OD) stainless steel cannulas for the fluidic interface. Stainless steel reinforcement tubes (OD = 635 µm) permitted interfacing with standard microdrive holders (Figure 1c,f). Up to eight probes connected to the same backend can be assembled into microdrives, with independent fluidic interfaces for each probe, enabling independent depth control of multiple fibers for broad-scale interrogation of neural activity (Extended Data Figure 1b).

The choice of materials and backend interface was informed by the need for thorough sterilization of the devices. The relatively high glass transition temperature (*T*_*g*_) of PEI allowed the fiber-based devices to be autoclaved without deforming, and encasing the backend electrode interface in medical epoxy protected the electrical components. The impedance magnitude (|*Z*|) of the integrated electrodes did not change significantly after autoclave sterilization (before |*Z*| = 223.9±36.6 kΩ, after |*Z*| = 206.7±22.9 kΩ; Figure 1g). Accurate infusion of nanoliter volumes at rates of 10–100 nL/min was achieved with 7.0±0.3 cm long fibers, with measured infusion rates matching those set by an infusion pump (Figure 1h). This resulted in accurate infusions, with mean absolute error (MAE) between the actual and expected infusion volume of 1.77 nL at 50 nL/min. Dynamic material analysis demonstrated that the stiffness of the fibers was 45.53 ± 1.14 N/m in the frequency range of respiration and heartbeat for NHPs (Figure 1i). This is significantly lower than the stiffness of a stainless steel cannula with comparable dimensions to preexisting multifunctional devices used in NHP research (OD = 203 µm, ID = 50 µm, stiffness = 180.99 ± 0.22 N/m).

### 2.2 Simultaneous intracortical drug delivery and neural activity recordings

To illustrate simultaneous pharmacological modulation and electrophysiological monitoring of neuronal activity in NHPs, we applied multifunctional fibers to record single unit and local field potential (LFP) activity while delivering gamma-amino butyric acid (GABA) in the premotor cortex. The waveforms from neuronal action potentials were recorded across the four electrodes, providing four observations of the same putative single unit (Figure 2a,b). Single units were isolated by clustering the principal components of waveforms across the four electrodes (Extended Data Figure 2). We found that the waveforms were stable between the baseline and recovery phases. The ability to record neural activity before, during, and after the infusions enabled tracking of putative single unit activity in response to intracranial GABA delivery (signal-to-noise ratio, SNR baseline = 3.08 [0.82, 26.15], SNR during = 5.42 [1.31, 16.27], SNR recovery = 9.61 [1.26, 24.42]; median [95% confidence intervals]; Figure 2c). In accordance with GABA’s role as an inhibitory neurotransmitter, we observed inhibition of single unit activity following five minute infusions of 100 mM GABA at 10 and 50 nL/min. Depending on the volume of GABA delivered (50 nL over 5 minutes, 250 nL over 5 minutes), we were able to cause inhibition of firing activity for both short (unit 2, 50 nL: 1.47 [1.28, 1.69] min) and long durations (unit 1, 250 nL: 10.13 [0.33, 12.38] min, unit 2, 250 nL: 26.27 [22.82, 28.43] min), followed by recovery of firing rates similar to or exceeding baseline rates (Figure 2d,e). Firing rates were estimated using a state-space point-process model (Methods, [7]). Downstates were defined as periods where the firing rate was below the minimum baseline firing rate for at least 10 consecutive seconds. 95% confidence intervals for firing rates, downstate times, and downstate durations were estimated with Monte Carlo (Methods).

**Figure 2:**
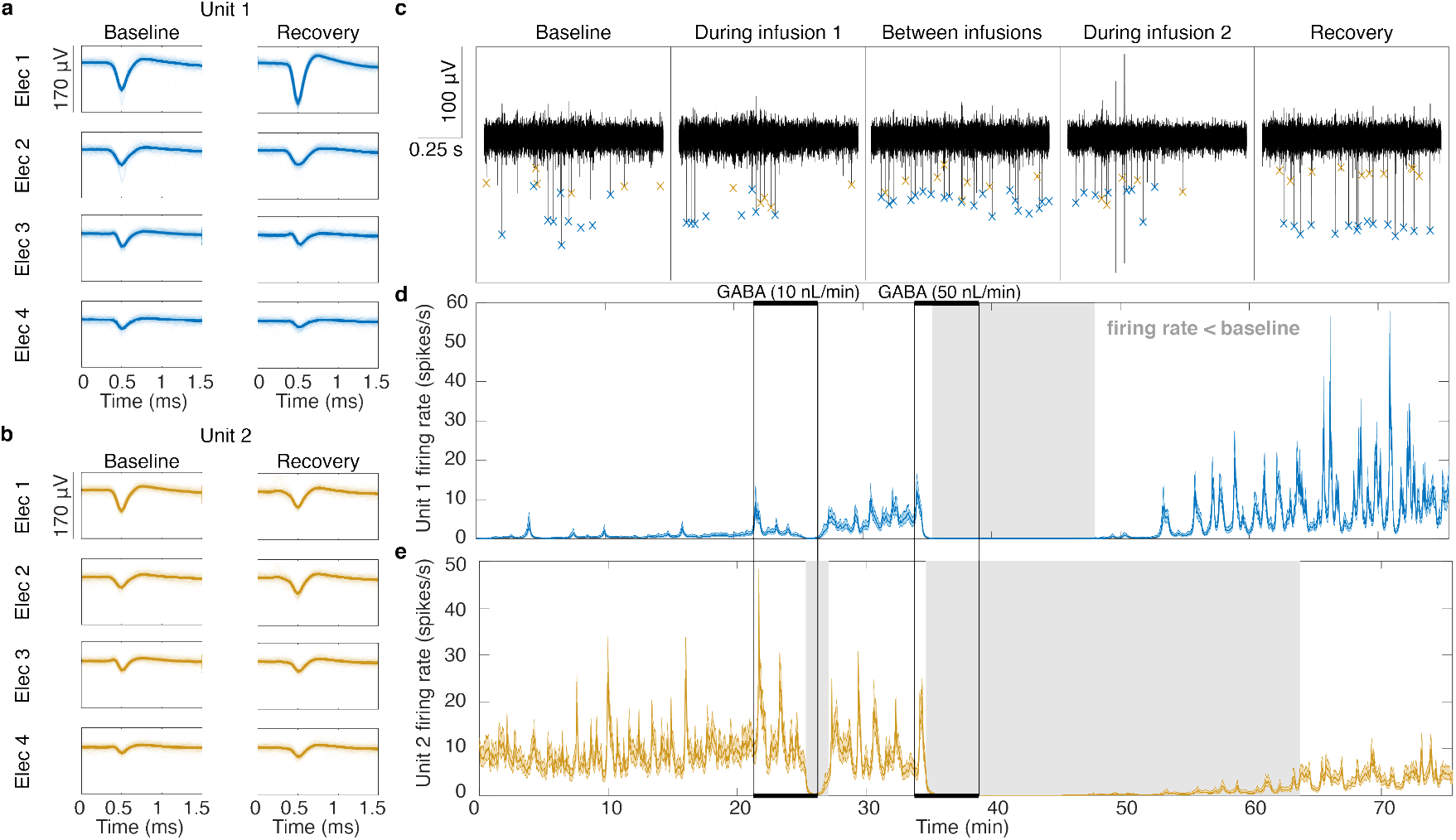
Multifunctional fibers record the inhibitory effect of intracortical 100 mM GABA delivery on single unit activity in real-time. **a–b**, Two units were recorded with the same probe, and their waveforms were observable across all four electrodes. The waveforms were stable in both shape and relative amplitude before and after 10 and 50 nL/min infusions of GABA. **c**, The noise floor of the recordings was not affected by intracortical GABA delivery, allowing for consistent identification of single unit waveforms before, during, and after two GABA infusions at 10 nL/min (infusion 1) and 50 nL/min (infusion 2). The blue x’s correspond to unit 1 and the yellow x’s to unit 2. **d–e**, The firing rate of single unit activity was modeled with a state-space point-process model (Methods). 95% confidence intervals for the estimated rate are shown with the blue and yellow shaded areas. Periods where the estimated firing rate was lower than the minimum baseline firing rate are indicated with grey rectangles. The infusion periods are indicated with black rectangles. **d**, The mean firing rate of unit 1 was 0.82 [0.78,0.89] spikes/second before the first GABA injection, decreased below baseline for 12.63 [8.10, 13.21] minutes following the 50 nl/min GABA injection, and then increased to 6.05 [5.30, 6.30] spikes/second. **e**, The mean firing rate of unit 2 was 9.10 [9.01, 9.34] spikes/second before the first GABA injection, decreased below baseline for 1.78 [1.38, 1.85] minutes following the 10 nl/min injection and 29.03 [26.25, 29.12] minutes following the 50 nl/min injection, and then recovered to 3.87 [3.48, 4.06] spikes/second.

Overall, 9 of 10 recorded neurons had significantly decreased mean firing rate in the 10 minutes following GABA infusions, with a subsequent increase in firing rate 20 minutes after the end of the infusion (Extended Data Figure 3). Thus, to compare neurophysiology over time, we define four periods: 1) the baseline period, before any GABA was delivered; 2) the infusion period(s), during intracranial GABA delivery; 3) the inhibition period, 10 minutes following GABA delivery; and 4) the recovery period, greater than 20 minutes after GABA delivery.

We also observed changes in the oscillatory structure of the LFP between the baseline and inhibition periods. Figure 3a shows the spectrogram of LFP recorded in the same session reported in Figure 2. We used the Fitting Oscillations and One Over F (FOOOF) algorithm to characterize LFP spectra [42]. The FOOOF algorithm models a given power spectral density (PSD, in decibels) as a sum of an aperiodic exponential decay component and periodic oscillation peaks (Methods). The aperiodic component has a broadband offset and rate of exponential decay which characterizes the decrease in power with increasing frequency, a common feature of neural spectra [42, 43]. We found dominant oscillatory peaks in three frequency ranges across four recordings: 11–17 Hz (mean center frequency, MCF: 13.46 [13.35, 13.61] Hz), 27–35 Hz (MCF: 29.73 [29.64, 29.82] Hz), and 47–62 Hz (MCF: 54.59 [54.03, 55.04] Hz; Figure 3c). Note that these ranges, estimated from the unsupervised FOOOF algorithm, correspond to the canonical alpha, beta, and gamma frequency ranges. We found that during the inhibition period following intracranial GABA infusion, there was a significant decrease in mean peak LFP power in the 11–17, 27–35 Hz and 47–62 Hz ranges (Δ_11*−*17_ = *−*0.58 [*−*0.97, *−*0.25] dB, Δ_27*−*35_ = *−*1.14 [*−*1.58, *−*0.72] dB, Δ_47*−*62_ = *−*0.54 [*−*0.88, *−*0.17] dB; Figure 3b–d), and an increase in mean peak power in the 35–47 and 70–100 Hz ranges (Δ_35*−*47_ = 0.23 [0.03, 0.45] dB, Δ_70*−*100_ = 0.15 [0.06, 0.25] dB; Figure 3c,d). We found an increase in both the offset and exponent of the aperiodic decay (Δ_*offset*_ = 0.04 [0.00, 0.08],Δ_*exponent*_ = 0.04 [0.03, 0.06]), which corresponded to a small broadband decrease in high-frequency LFP power following cortical GABA infusions.

**Figure 3:**
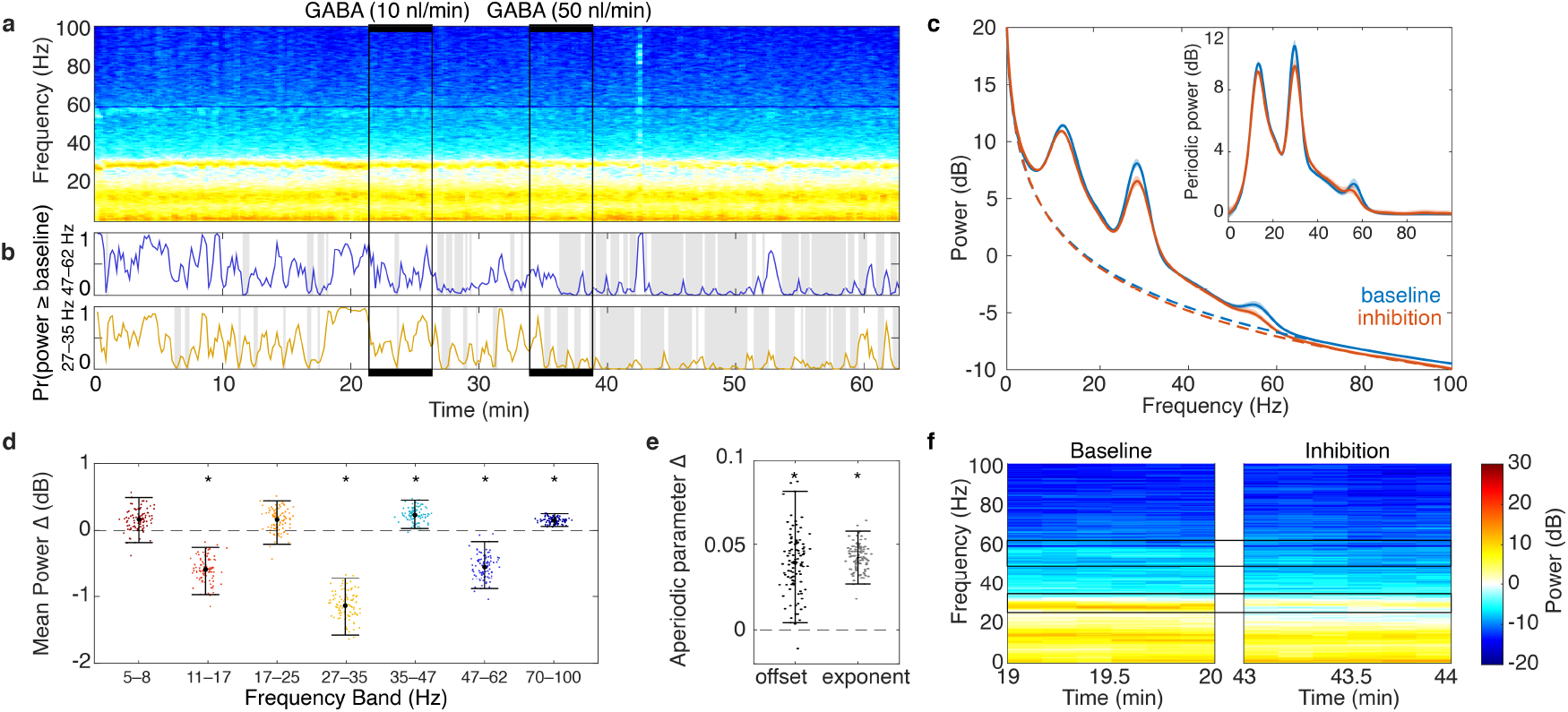
Intracortical 100 mM GABA delivery alters the oscillatory structure of local field potential (LFP) activity. **a**, A multitaper spectrogram was calculated from the LFP activity recorded during the same experiment presented in Figure 2. **b**, The probability that peak power in the 27–35 Hz and 47–62 Hz bands is greater than or equal to the peak power before GABA was delivered (*Pr*(power *≥* baseline)) are plotted in yellow and blue, respectively. Periods where this probability is less than 0.1 are indicated with the grey boxes. After the 50 nL/min infusion, there is sustained suppression of oscillatory peak power in both the 27–35 and 47–62 Hz bands. **c**, The mean and 95% confidence intervals of 100 bootstrap samples of the mean spectral power (dB) calculated from 10-minute periods before and after GABA are indicated in blue and red, respectively. The dotted lines show the mean aperiodic component across all bootstrap samples before and after GABA. The inset shows the mean and 95% confidence intervals for the spectral power minus the aperiodic component (Periodic power). **d**, 100 bootstrap estimates of the change in mean peak power in the 10 minutes before the first GABA infusion vs. the 10 minutes after the last GABA infusion; median and 95% confidence intervals are denoted with black dots and black horizontal lines, respectively. Asterisks indicate bands where the 95% confidence intervals of the mean power difference do not include zero. **e**, The mean estimated offset and exponent of the aperiodic component were increased following GABA infusion. Together, the reductions in peak power and increase in the exponent of aperiodic decay result in reduced power following local GABA delivery. **f**, Side-by-side comparison of periods of the spectrogram calculated from before the first GABA infusion and after the second GABA infusion show reduced power, most dominant in the 27–35 and 47–62 Hz bands.

To control for the effect of the infusion itself on neurophysiology recordings, we recorded neural activity in response to the GABA vehicle, saline (Extended Data Figure 4). Saline did not cause a significant decrease in firing rate, although there was a small reduction in spike amplitude (SNR baseline = 2.80 [0.92, 6.25], SNR during infusion = 2.49 [1.06, 5.14], SNR after infusion = 2.83 [1.13, 6.67]), which may stem from minor tissue deflection during infusion. The changes in the 11–17, 27–35, and 47–62 Hz frequency bands following saline infusions were significantly smaller than with the GABA infusions (Extended Data Figure 4e).

### 2.3 Modulation of cognitive task encoding

We hypothesized that, in addition to locally reducing firing rates and altering LFP oscillations, intracranial GABA infusions may locally alter cognitive encoding in the premotor cortex (Figure 4a). To test this hypothesis, we recorded neural activity while the NHP performed a delayed match-to-sample (DMS) working memory task [38, 44, 45] (Figure 4b,c). In each trial, after fixating on the center of a screen, the NHP was presented with one of three images for 1 second (sample phase). The image then disappeared for 1 second (delay phase). Then, two images appeared on the screen in the top right and bottom left corners and the NHP saccaded to one of the images (match phase). If the NHP selected the image presented during the sample phase, they were rewarded with a juice pulse (reward phase). If not, the trial ended with no reward.

**Figure 4:**
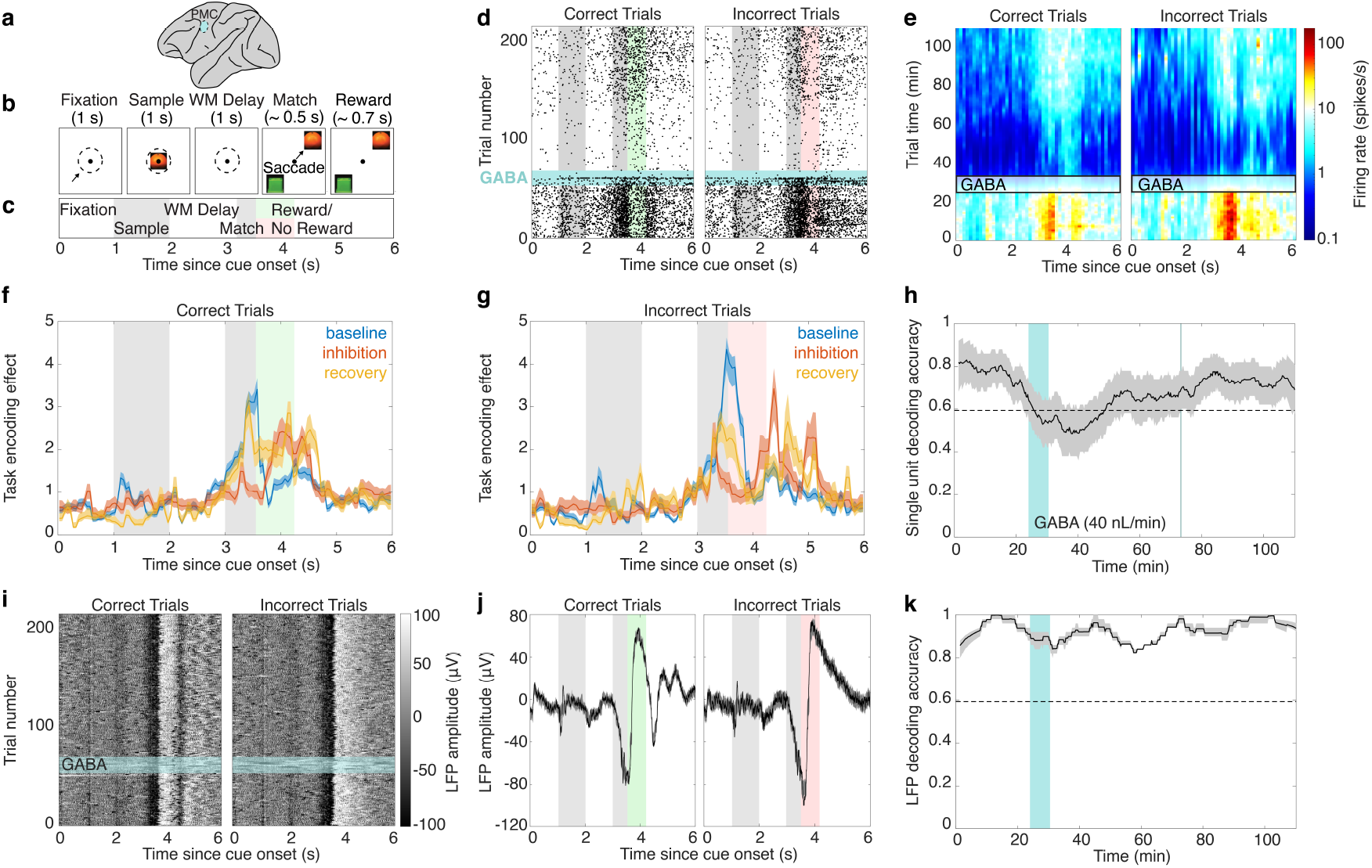
Intracranial GABA infusions alter neural encoding during a working memory task. **a**, Multifunctional fibers were implanted in the premotor cortex. **b**, The delayed match-to-sample (DMS) working memory (WM) task involves 1 second fixation, sample presentation, and delay phases. The NHP then saccades to the choice that matches the sample. If the choice is correct, juice is dispensed. If the choice is wrong, no juice is dispensed. **c**, In panels d, f, g, and j, we use shaded bars to indicate each task phase. The shaded bar between 1–2 seconds indicates the sample phase, the shaded bar between 3–3.5 seconds indicates the match phase, and the colored bar (green for correct trials, red for incorrect trials) from 3.5–4.2 seconds indicates the average timing of the reward phase during correct trials. **d**, Raster plots of single unit activity during correct and incorrect trials. The first vertical grey bar marks the phase of sample presentation, the second grey bar marks the time when the choices appear on the screen and a decision is made, and the green or red bars marks the phase when juice is dispensed during correct trials. The horizontal blue bar marks the trials during which GABA (100 mM, 40 nL/min) was infused. 224 correct trials and 224 incorrect trials from one session are shown here. **e**, State-space generalized linear models (SS-GLMs) were used to estimate the firing rate within and across trials. The horizontal box indicates when GABA was infused. Task phase was a significant predictor of firing rate (p *<* 0.001, likelihood ratio test). There was a significant reduction in firing rate following GABA infusion (baseline firing rate = 10.90 [10.81, 10.99] spikes/s, inhibition period = 0.86 [0.85, 0.88] spikes/s), that increased during the recovery period (2.34 [2.33, 2.35] spikes/s). **f**, After factoring out the trial average firing rate, the SS-GLM provides an estimate of the task encoding effect over time (Methods). The task encoding effect during correct trials varied significantly between the baseline (blue), inhibition (red), and recovery (yellow) periods. **g**, The task encoding effect during incorrect trials also varied significantly between the baseline (blue), inhibition (red), and recovery (yellow) periods. **h**, As task encoding varied throughout the experiment, so did correct vs. incorrect trial decoding accuracy (black line = mean decoding accuracy across sliding windows of 50 consecutive trials, grey shading = 95% confidence intervals). The dotted line indicates the null decoding accuracy (Methods), the vertical blue bar indicates when GABA was infused, and the green line indicates the first time after GABA where the 95% confidence intervals for the decoding accuracy were greater than the null decoder. **i**, Local field potential (LFP) amplitude across trials. The same trials plotted in panel b are shown here. Dark shades indicate low LFP amplitude, while light shades indicate high LFP amplitude. The horizontal blue bar marks the trials during which GABA was infused. **j**, An autoregressive model with trial covariates characterizes LFP amplitude within trials. The black line indicates the mean LFP amplitude across all trials, and the 95% confidence intervals for the mean LFP amplitude are indicated in grey. **k**, Correct vs. incorrect trial decoding accuracy using LFP observations was above the null decoding accuracy throughout the experiment (black line = mean decoding accuracy across sliding windows of 50 consecutive trials, grey shading = 95% confidence intervals). The dotted line indicates the null decoding accuracy (Methods), and the vertical blue bar indicates when GABA was infused.

Consistent with previous studies of working memory [38, 39], we observed that the single unit firing rate in the premotor cortex varied consistently within trials (Figure 4d). To characterize this activity and how it changed during and following GABA delivery, we modeled the time-varying relationship between spiking activity and trial phase using a state-space generalized linear model (SS-GLM) [8, 46]. In this analysis, we separated the trial into 50 evenly spaced subphases and investigated how the firing rate varied between each subphase. We modeled single unit activity as a point process, where the instantaneous firing rate varied as a function of the firing history of the single unit and the trial subphase (Methods). This captured firing dynamics within a trial that were driven by innate properties of the neuron, such as refractory period, and those that were driven by task encoding. Each trial had a unique set of trial subphase coefficients, which were modeled as latent state variables that vary smoothly between sequential trials. The latent state process characterized how firing rate and task encoding changed across trials. Overall, the SS-GLM provided a unified model for characterizing changes in firing rate within and across trials. Trials were ordered sequentially, and the model was estimated from odd (training) trials, while goodness of fit was assessed from even (test) trials (Extended Data Figure 5). We found that trial subphase was a significant predictor of firing rate for 11/12 single units recorded across 6 recording sessions (Likelihood Ratio test, *p <* 0.05, Methods; Extended Data Figure 6).

After identifying that the firing rate of a unit varied between trial subphases, we investigated if task evoked activity varied between trial variants (left vs. right saccades, sample ID, choice ID, correct vs. incorrect trials). We separately estimated models from training trials belonging to each trial variant, and then compared the difference in task evoked activity between each variant (Extended Data Figure 9). Figure 4d–g presents a unit for which task evoked activity varied most significantly between correct vs. incorrect trials. GABA delivery reduced the unit’s firing rate and also restructured its task encoding (Figure 4d). The SS-GLM provided a quantitative characterization of these results (Figure 4e). Together, the multifunctional fibers and SS-GLM provided the opportunity to investigate how single unit task encoding changed following local inhibition. By factoring out the estimated trial-average firing rate, the variation in firing rate due to the task (the task encoding effect) could be isolated and compared between the baseline, inhibition, and recovery periods (Figure 4f,g). Prior to GABA infusion, this unit had a slight increase in firing rate at the beginning of the sample phase in both correct and incorrect trials. Then, the firing rate gradually increased from the delay phase to the time that a decision was made at the end of the match period. During correct trials, the firing rate rapidly dropped when juice was dispensed. During incorrect trials, the firing rate continued to rise before decreasing in the middle of the phase when juice would have been dispensed if the trials were correct. Thus, before GABA, this unit encoded information across multiple trial phases, including the presence or absence of reward. We found that local GABA infusions altered the task encoding effect: 4.0 minutes into the GABA infusion, there was a significant reduction in firing rate across a majority of trial subphases, but the inhibitory effect was not uniform across subphases. While this neuron had the highest firing rate during the match phase before GABA, in the inhibition period, its firing rate was highest during and following the reward phase. Further, the increase in firing rate at the beginning of the sample phase was lost during the inhibition period. As the effect of the GABA infusion wore off, there was an increase in firing rate during the match phase, but the increase in firing at the beginning of the sample phase did not return. As a result, 65–80 minutes after GABA, the 95% confidence intervals of the task encoding effect had 55.0% and 62.5% overlap with the baseline task encoding effect for correct and incorrect trials respectively, indicating incomplete recovery of task encoded activity.

We investigated how changes in task encoding caused by local GABA infusions affected the ability to decode behavioral information from single unit activity (Figure 4h). We decoded correct vs. incorrect trials by comparing the deviance of the data observed during test trials from two SS-GLMs estimated from correct or incorrect training trials. Before GABA, it was possible to decode whether test trials were correct or incorrect with 80 [72, 87]% accuracy from the activity of one single unit alone, which is significantly above the null decoding accuracy of 59% (Methods). As the task encoded activity of the neuron shifted in response to GABA, the ability to reliably decode correct vs. incorrect trials dropped below the null decoding accuracy (decoding accuracy during the inhibition phase = 49 [38, 61]%). The lower 95% confidence interval of the decoding accuracy first passed above the null decoding accuracy 43.3 minutes after the end of the GABA infusion, when the trial average firing rate was 16.1 [7.4, 35.8]% of the baseline levels and task encoding remained significantly different from baseline. Thus, despite persistent changes in the task encoded activity, there was sufficient information encoded in the signal to distinguish correct vs. incorrect trials above chance.

We found that local GABA delivery had an opposite effect on encoding of task-related LFP activity. First, task related LFP activity was relatively stable throughout the experiment (Figure 4i), despite changes in the oscillatory structure of the LFP (Figure 3). We modeled the relationship between LFP amplitude and trial phase using an autoregressive model with trial phase covariates. The mean and 95% confidence intervals of the event related potentials for correct and incorrect trials are shown in Figure 4j. We found that LFP was a reliable marker of whether a trial was correct or incorrect (94 [93, 96]% accuracy before GABA; Figure 4k). In contrast to single unit activity, the decoding accuracy from LFP was stable during and following local GABA delivery (89 [86, 92]% and 89 [86, 91]% accuracy during the infusion and inhibition phases, respectively; Extended Data Figure 7).

To assess if changes in neurophysiology caused by GABA affected behavioral performance, we compared task accuracy before and after GABA. We observed a significant increase in task accuracy in one of the GABA infusion sessions, but this effect was not repeated across sessions (Extended Data Figure 8).

### 2.4 Intrastriatal modulation of neural activity and cognitive task encoding

We leveraged the unrestricted length of the fibers to record the effect of intrastriatal GABA delivery on single unit and LFP activity in the putamen (Figure 5a), a subcortical structure with multiple functions, including processing reward [40, 41]. Due to the proximity of the putamen to important regulatory centers, we infused a smaller volume of GABA as compared to the cortical recordings (putamen: 56 nL total; premotor cortex: 222.8 ± 27.8 nL total). As in the cortical recordings, the same single unit was recorded before and after GABA delivery (Figure 5b). This unit’s task encoded activity varied most between correct and incorrect trials, particularly during and immediately following the reward phase (Figure 5c–f). As in the premotor cortex analysis, the SS-GLM characterized how task encoded activity varied following local GABA delivery (Figure 5d). There was a decrease in trial-average firing rate during the GABA infusion (firing rate in the trial preceding GABA = 8.27 [7.59, 8.99] spikes/s, firing rate in the trial following GABA = 4.76 [4.36, 5.22] spikes/s), followed by an increase in firing rate in specific trial phases (Figure 5d). Figure 5e,f characterize how the task encoding effect varied during the baseline, post-GABA, and recovery period. The most significant changes included an increase in task modulation following the reward phase in incorrect trials during the recovery period. This manifested as a greater difference between correct and incorrect trials and an increase in single unit decoding accuracy during the recovery period.

**Figure 5:**
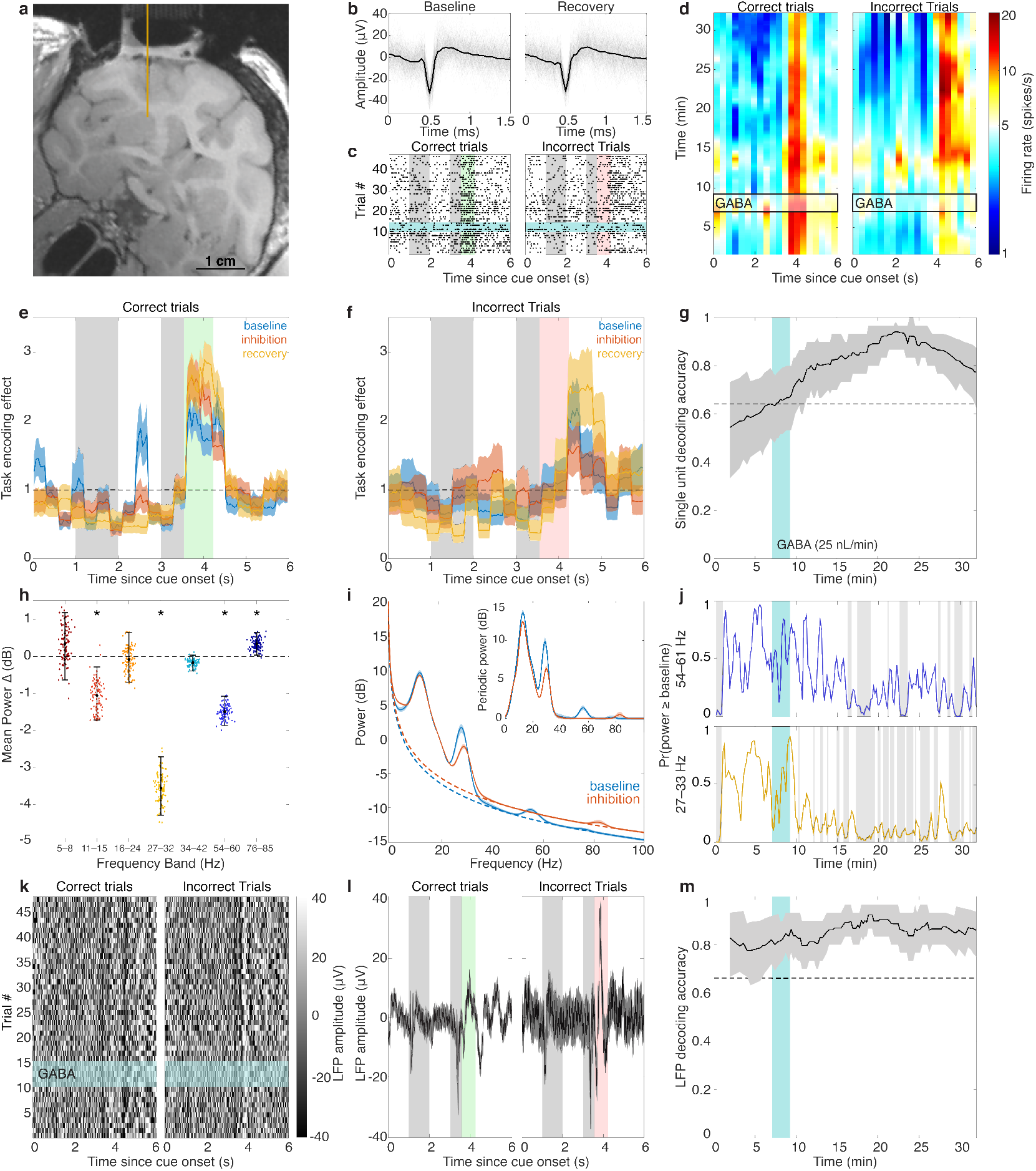
Multifunctional fibers in the putamen record the effect of intrastriatal GABA infusion on single unit and LFP reward encoding. **a**, MRI of the NHP subject used in this study. The yellow line indicates the implantation trajectory of the multifunctional fiber probe to the putamen. **b**, The waveform of single unit activity was stable before and after GABA (25 nL/min) was delivered. **c**, Raster plot of single unit activity during correct and incorrect trials of the same task illustrated in Figure 4 (horizontal blue bar: GABA infusion). **d**, Single unit firing rate varied within and between trials. Task phase was a significant predictor of firing rate (*p <* 0.001, likelihood ratio test). There was a significant reduction in firing rate during the GABA infusion (firing rate in the trial preceding GABA = 8.27 [7.59, 8.99] spikes/s, firing rate in the trial following GABA = 4.76 [4.36, 5.22] spikes/s). **e–f**, The task encoding effect during correct (**e**) and incorrect (**f**) trials varied significantly between the baseline (blue), inhibition (red), and recovery (yellow) periods. **g**, Decoding accuracy for correct vs. incorrect trials was above the null decoding accuracy after GABA was infused. (black line: mean decoding accuracy across sliding windows of 30 consecutive trials, grey shading: 95% confidence intervals, dotted line: null decoding accuracy (Methods), vertical blue bar: GABA infusion) **h**, LFP oscillations were altered by GABA infusions—the change in mean peak power between 10 minutes before and after GABA infusion. Asterisks indicate bands where the 95% confidence intervals of the mean power difference do not include zero. **i**, The mean and 95% confidence intervals of 100 bootstrap samples of the mean spectral power (dB) before and after GABA are indicated in blue and red, respectively. The dotted lines show the mean aperiodic component across all bootstrap samples before and after GABA. The inset shows the mean and 95% confidence intervals for the spectral power minus the aperiodic component (Periodic power). **j**, The probability that peak power in the 27–32 Hz and 54–60 Hz bands was greater than or equal to the peak power before GABA was delivered (*Pr*(power*≥* baseline)) are plotted in yellow and blue, respectively. Periods where this probability was less than 0.1 are indicated with the grey boxes. **k**, LFP amplitude across trials. Dark shades indicate low LFP amplitude, while light shades indicate high LFP amplitude. The horizontal blue bar marks the trials during which GABA was infused. **l**, An autoregressive model with trial covariates characterized LFP amplitude within trials. The black line indicates the mean LFP amplitude across all trials, and the 95% confidence intervals for the mean LFP amplitude are indicated in grey. **m**, Correct vs. incorrect trial decoding accuracy using LFP observations was above the null decoding accuracy throughout the experiment.

Similar to the premotor cortex recordings, there was a modulation of LFP oscillatory content following GABA delivery. Using the FOOOF algorithm, we found there was a significant decrease in the 11– 15, 27–32, and 54–60 Hz bands (Δ_11*−*15_ = *−*1.05 [*−*1.72, *−*0.29] dB, Δ_27*−*32_ = *−*3.56 [*−*4.29, *−*2.71] dB, Δ_54*−*60_ = *−*1.5 [*−*1.72, *−*0.29] dB), and a significant increase in the 76–85 Hz band (Δ_76*−*85_ = 0.33 [0.03, 0.63] dB; Figure 5h,i). In contrast to the premotor cortex, there was a decrease in the rate of exponential decay (Δ_*exponent*_ = *−*0.04 [*−*0.06, *−*0.02]), resulting in elevated broadband high frequency activity (Figure 5i). Figure 5j shows the time course of the change in LFP oscillatory power in the 27–32 and 54–60 Hz bands.

As in the premotor cortex, we found that despite these changes in the oscillatory structure, task-evoked potentials were relatively stable throughout the experiment (Figure 5k). Task evoked LFP activity varied significantly between correct and incorrect trials (Figure 5l), which enabled accurate decoding of correct and incorrect trials from LFP throughout the experiment (Figure 5m, Extended Data Figure 7). We did not find any significant changes in task performance compared to pre-GABA task accuracy (Extended Data Figure 8).

## 3 Discussion

Multifunctional fibers enabled detailed investigation of the effect of local inhibition on cognitive neural encoding in an awake, behaving macaque. Our probe design was informed by recent research in flexible neurotechnology, which has emphasized the importance of minimizing stiffness of intracranial devices [25–29, 34–37]. By minimizing the fiber diameter and using flexible PEI, we achieved autoclavable devices with significantly lower stiffness than state-of-the-art probes used for intracranial drug delivery experiments in NHPs (Figure 1). We established that our devices are able to record neural activity before, during, and after intracranial GABA delivery. We then investigated the effect of intracranial GABA delivery on single unit and local field potential activity in the premotor cortex and putamen while the macaque performed a working memory task. We found that, while GABA had the anticipated inhibitory effect on firing rates, single unit and LFP activity encoded task-related information even while firing rates were suppressed. By locally modulating neural excitability, we probed how information encoded in intact neural circuits may be remodeled in the presence of neuromodulatory compounds.

*In vivo* neuromodulation affects neural dynamics across time scales ranging from seconds to hours. To characterize this broad dynamic range, we employed several time series analysis methods. First, we characterized the change in neuronal firing rate during and after local GABA infusion by estimating a state-space point-process model of single unit activity (Figure 2). Then, to gain more detailed insight into how local GABA infusion affected neural computations, we used a state-space generalized linear model to characterize variations in firing rate within and across trials of the working memory task (Figure 4).

Modeling task encoding over time with the SS-GLM enabled us to not only capture broad changes in firing rate due to GABA delivery, but also how single unit encoding of cognitive task information evolved throughout a period of temporary inhibition. These results emphasize that cognitive encoding by single units is dynamic [8, 47], and can be altered by changes in local concentrations of neurotransmitters such as GABA.

We found that local drug delivery caused sustained changes in the oscillatory structure of LFP (Figure 3). Local inhibition resulted in significant reduction of oscillatory power in the prominent alpha, beta, and gamma frequency bands, each of which have been associated with cognitive function, including working memory [4, 38, 45]. Despite changes in oscillatory content, task evoked potentials provided a reliable signal for decoding task information, even during periods of near complete spike inhibition. Task evoked LFP activity is associated with both local spiking and coordinated synaptic activity [43]. We hypothesize that the primary contribution to task evoked potentials was synaptic input from neural populations unaffected by local GABA infusions. Using multifunctional probes to investigate the relative contribution of local spiking and coordinated synaptic input would enable further exploration of this hypothesis.

Finally, while previous studies of multifunctional neurotechnology have discussed the value of studying deep brain neurophysiology [24], simultaneous electrophysiology and drug delivery in deep brain structures remains rare. To our knowledge, there has only been one previous study of simultaneous iontophoretic drug delivery and electrophysiology in the striatum of an awake, behaving macaque [48]. However, there has been considerable interest in refining drug delivery to deep brain structures for therapeutic purposes [15, 49–51], although physiological measures of the effect of deep brain drug delivery have been limited. The dearth of studies investigating the electrophysiological effects of drug delivery in deep brain structures may be related in part to safety concerns, particularly for the implantation of large, stiff, or fragile devices near critical vasculature [52]. Our devices address the limitation of existing stainless steel technology by minimizing diameter and stiffness (Figure 1) [22–24]. Compared to glass iontophoresis probes, our devices have greater flexural strength and are thus less susceptible to fracture.

While our present study was limited to a single subject, our devices were designed to be compatible with commercially available microdrive systems and standard headstages, which will facilitate dissemination of this technology to the NHP neuroscience community. Future device development will include extending the device functionality along the length of the fiber to increase electrode count and incorporating functionalities routinely available for rodent research [34, 35, 37], such as optical channels for optogenetics. We developed multifunctional fibers for modulating neural activity in NHPs and applied state-space models designed to track neural dynamics over broad timescales, which together uncover a path toward detailed investigations into the effects of neuromodulation on intact neural circuits during complex cognitive tasks.

## 4 Methods

### 4.1 Device Fabrication

#### 4.1.1 Fiber

We fabricated multifunctional fibers with four tungsten microelectrodes (25 µm diameter), two microfluidic channels (40 × 50 µm^2^), and a poly(etherimide) (PEI, glass transition temperature, *Tg* = 217°C) cladding. Each fiber had a diameter of 150–200 µm. The fibers were fabricated with the thermal drawing process (Figure 1), which has been previously described in detail [34, 35, 53]. Briefly, thermal drawing begins by first fabricating a fiber preform. In this case, we milled four 1 mm × 1 mm electrode channels and two 2 mm × 2.5 mm fluidic channels in a PEI rod (diameter = 6.35 mm, length = 20 cm, McMaster-Carr, #8686K71). We then rolled 25 µm thick PEI film (McMaster-Carr, #7576K15) around the rod to achieve a preform with outer diameter of 7.5 mm. We consolidated the outer layers of polymer with the inner rod by heating the preform at 250°C for 20 minutes. The preform was then placed in a cylindrical thermal drawing oven heated to 320°C. Electrodes were incorporated into the fiber using a process called convergence. Four spools of tungsten microwire were oriented above the thermal drawing oven and the ends of the wire were fed into the electrode channels of the preform. Heating the preform caused the polymer at the center of the oven to flow. As the polymer flowed, initially gravity caused the end of the preform to fall downwards, resulting in a section of reduced diameter and increased length. When the lower end of the preform reached a capstan situated below the oven, the lower portion of the preform was cut away and the necking point was fed into the capstan. The capstan velocity (*v*_capstan_) was progressively increased while the preform was fed further into the oven (*v*_downfeed_). The ratio between *v*_capstan_ and *v*_downfeed_ determines the ratio between the cross-sectional area of the preform and fiber:

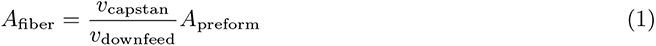

Here, we used *v*_capstan_ = 0.8 m/min. and *v*_downfeed_ = 0.5 mm/min for a ratio of 1:40 between the fiber and preform diameters. As the fiber diameter decreased, the electrode channels reduced in size. When the channels decreased to the size of wire, they constricted around the wire and pulled them into the fiber at the same rate that the fiber was drawn.

#### 4.1.2 Back-end Interface

The fiber was then cut into 10 cm sections and connected to a custom back-end interface. Our back-end interfaces are highly modular, and can be modified to connect one to eight fibers to the same headstage connector (Blackrock Neurotech, Cereplex M). Each individual fiber has a stainless steel support tube (432 µm inner diameter (ID), 635 µm outer diameter (OD), McMaster-Carr, #75165A685) and microfluidic connector tube (304 µm ID, 457 µm OD, McMaster-Carr, #75165A556). The tungsten electrodes were bonded to 50 µm stainless steel microwires with silver epoxy (Epo-Tek, #H20E), which were then bonded to electrode connectors (Omnetics Connector Corporation, A79026, A79014). In a single fiber configuration (Figure 1), the electrode connector was epoxied to the support tube of the fiber with medical epoxy (Loctite EA M-31CL).

#### 4.1.3 Sterilization

All fiber and backend materials were selected to allow for autoclave sterilization of the devices. The connected devices were steam autoclaved at 121°C for 45 minutes. After assembly in the cranial grids (Section 4.2, Extended Data Figure 1), the tips of the fibers were soaked in chlorhexidine diacetate (Novalsan) and rinsed with sterile water.

#### 4.1.4 Characterization

Electrical impedance spectroscopy was performed before and after sterilization using an LCR meter (HP4284A, Agilent Technologies) with sinusoidal input voltages (20 mV, 0.1–100 kHz). The infusion rate through microfluidic channels was characterized by injecting PBS saline at 10–100 nL/min through the fiber (NanoFil Syringe and UMP3 Syringe pump, World Precision Instruments) and imaging the rate of flow through a capillary attached to the outlet of the fiber. The stiffness of the fibers (*n* = 3) compared to a stainless steel cannula was characterized with dynamic material analysis (Q800, TA Instruments). DMA was performed in single-cantilever mode, using 17.5 mm long samples and 50 µm vertical deflections.

### 4.2 Animal Experiments

#### 4.2.1 Recordings

All surgical and animal care procedures were approved by the Massachusetts Institute of Technology (MIT)’s Committee on Animal Care and were conducted in accordance with the guidelines of the National Institute of Health and MIT’s Department of Comparative Medicine (Protocol 0619-035-22). Experiments were conducted with one female macaque (*macaca mulatta*, age = 21 years, weight = 9.3 kg). Probes were connected to a NAN-drive micromanipulator (NAN Instruments, NAN C) via the probe support tube and assembled in a cortical chamber (Extended Data Figure 1). GABA or saline was prefilled in all infusion lines. While the NHP was seated in a primate chair, the cortical chamber was placed in a custom-made cephalic chamber implant (surgical procedure described in [45]). A 24-gauge guide tube was lowered to pierce the dura. Probes were then lowered at a rate of 0.05 mm/s while LFP and spiking activity was continuously monitored. Probes implanted in the premotor cortex were approximately 1.5 mm below the cortical surface, and probes implanted in the putamen were 12.4 mm below the cortical surface. Probes then settled for at least 60 minutes. Baseline recordings were collected for 7–30 minutes. Then, either GABA or saline was injected (Parker Picospritzer III). For all cortical infusions except one low-rate GABA infusion (Figure 2), the target infusion rate was 50 nL/min. Infusion rate was set by adjusting the infusion pressure and the exact rate of infusion was measured by tracking the velocity of the meniscus through the drug tubing (Molex, Polymicro fused silica capillary tubing, ID = 100 µm). Actual cortical infusion rates were 59.0 ± 9.1 nL/min. The putamen GABA infusion rate was 25 nL/min. Total infusion volumes were 222.8 ± 27.8 nL for the cortical experiments and 56 nL for the putamen experiment. There were 4 premotor cortex recordings with GABA microinfusions, 3 premotor cortex recordings with saline microinfusions, and 1 putamen recording with a GABA microinfusion. To compare neurophysiology over the course of GABA recordings, we define four periods: 1) the baseline period, before any GABA was delivered; 2) the infusion period(s), during intracranial GABA delivery; 3) the inhibition period, 10 minutes following GABA delivery; and 4) the recovery period, greater than 20 minutes after GABA delivery (Results Section 2.2).

#### 4.2.2 Delayed-Match-to-Sample Task

Recordings were collected while the macaque performed a working memory task (Figure 4). The macaque sat in a primate chair inside a behavioral testing booth, 50 cm away from an LCD monitor (ASUS, Taiwan). Using positive reinforcement, we trained the macaque to perform a visual delayed-match-to-sample task. Eye movements were tracked with a video-based eyetracker (SR-Research, EyeLink 1000 Plus). In each trial, the macaque fixated on a gray point in the center of the screen (2 visual degrees radius) for 1 second. One of three cue images was briefly shown (1 second) and held in short term memory over a delay period (1 second). Then two images were presented (one matching image and one distractor image) at 8 degrees eccentricity. Images were randomly presented at either the upper right or lower left quadrant of the screen. The macaque then saccaded to one of the two images. If the selected image matched the image presented in the sample phase, the macaque received a juice reward.

### 4.3 Data Analysis and Statistics

#### 4.3.1 Confidence Interval Notation

For variables where the quantity is expected to follow a Gaussian distribution, we report the mean ± standard error of the mean. For other quantities, including those computed through bootstrap and Monte Carlo estimation (Sections 4.3.2 and 4.3.3) we report the median [2.5^*th*^ percentile, 97.5^*th*^ percentile].

#### 4.3.2 Single Unit Analysis

##### 4.3.2.1 Signal-to-Noise Ratio of Spike Recordings

Electrophysiology data was collected at 30 kHz and separated into high- and low-frequency data. High-frequency data was filtered between 0.25–10 kHz. Spike waveforms were extracted by identifying negative threshold crossings (*y*_*t*_ *< −*3.5*σ*, where *y*_*t*_ is the amplitude and *σ* is the standard deviation of the 0.5– 10 kHz filtered data) and extracting data from 0.5 ms before and 2 ms after each threshold crossing. Threshold crossings were measured across all four electrodes, and waveforms from all four electrodes were extracted for a threshold crossing on any electrode. Spikes were sorted into single unit activity by k-means clustering on the principle components computed from 2.5 ms spike waveforms on four electrodes. The signal to noise ratio (SNR) of the spike recordings was estimated by randomly sampling spike waveforms from each experimental period (baseline, during infusions, and recovery) and comparing the mean-squared amplitude of the waveform to an equal duration of randomly sampled spike-absent data from the same phases,

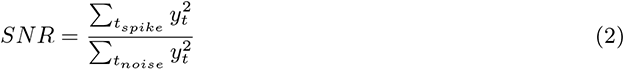

where *y*_*t*_ is the amplitude of the filtered data, *t*_*spike*_ denotes time points during each spike waveform sample, and *t*_*noise*_ denotes time points during each noise sample. We report the median and 95% confidence intervals of the SNR across 1000 spike samples.

##### 4.3.2.2 Estimating Firing Rate with Point Process Models

To characterize the firing rate of single units, we model single unit activity as a point process [6–8, 46]. For a given window of time, the probability of an action potential can be characterized with an instantaneous event rate parameter *λ*_*t*_. The instantaneous firing rate of a neuron is dependent on inherent properties, like refractory periods and interconnected circuit dynamics, as well as external factors, like behavioral stimuli. Thus, we model single unit activity with instantaneous spike rate conditioned on the underlying state of the unit using a point process model. The instantaneous rate of single unit activity is characterized by the conditional intensity function,

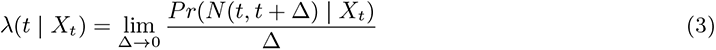

where *X*_*t*_ refers to the state of the neuron at time *t*, which is influenced by spike history and other modulating factors. *N* (*t, t*+Δ) denotes the number of spikes in the interval (*t, t*+Δ], and *Pr*(*N* (*t, t*+Δ) | *X*_*t*_) is the probability of *N* spikes given the state at time *t*. The conditional intensity function *λ*(*t* | *X*_*t*_) characterizes a point process model of spiking activity, where inter-spike intervals are dependent on the underlying state of the unit.

We employ two methods for characterizing single unit firing rate over time. The first is a state-space point process model [7], where the sources of firing rate variation (e.g., spike history, encoding of behavioral information, drug modulation) are characterized by a state variable *x*_*t*_, such that

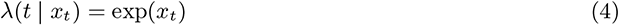

We define the state model as a first order Gaussian process model,

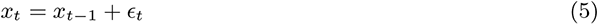

where *ϵ*_*t*_ is a Gaussian random variable with zero mean and variance *σ*^2^. Using this approach, we characterize how firing rate varies before, during, and after local GABA infusion. We estimated 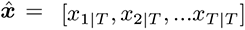, and 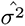 using the Expectation-Maximization (EM) algorithm [7], where *x*_*t*|*T*_ is the estimate of *x*_*t*_ given *N*_1:*T*_. 95% confidence intervals for *λ*(*t* | *x*_*t*|*T*_) were estimated with Monte Carlo [7]. From the estimated model, we computed 95% confidence intervals for the onset and offset of periods of significant inhibition following local GABA infusions as follows. Using the same algorithm used to estimate confidence intervals for *λ*(*t* | *x*_*t*|*T*_) [7], we generated *I* = 1000 realizations of 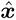. From each 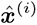, we computed the minimum firing rate before the first GABA infusion, 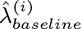. We then identified the time points 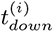 where 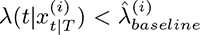 for at least 10 s. For a given downstate, 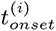 and 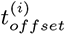 are the start and end of the downstate obtained from the *i*^*th*^ Monte Carlo realization of 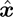. 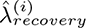 is defined as the mean firing rate computed from 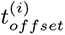 to *T*. The 95% confidence intervals for 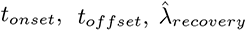, downstate duration, and average firing rate during a given segment (e.g., before the first GABA infusion) are the 2.5^*th*^ and 97.5^*th*^ percentiles of each quantity computed across *I* realizations of 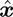..

The second method to model firing rate characterized both within trial and across trial dynamics. We use a state-space generalized linear model (SS-GLM) approach to characterize the dynamic relationship between instantaneous firing rate, trial phase, and spike history [8, 46]. We hypothesized that local inhibition of activity may affect both the rate and task encoding. Thus, with the SS-GLM approach, we assume that a state variable modulates the relationship between instantaneous rate and task encoding. Section 4.2 describes the task, which occurs in 6 second trials. We model the relationship between instantaneous firing rate and trial phase across *K* trials by considering *R* evenly spaced within-trial subphases. The trial subphase covariate, *S*_*t*_ is a set of *R* unit pulse functions, where *s*_*r,t*_ = 1 if time *t* is within trial subphase *r*. For the history covariate of the *k*^*′*^*th* trial, *H*_*k,t*_, we consider *J* = 9 non-overlapping history bins to capture the time-scales of common temporal dependencies between action potentials: 0–1 ms, 1–5 ms, 5–0 ms, 10–15 ms, 15–20 ms, 20–30 ms, 30–40 ms, 40–50 ms, and 50–100 ms (Extended Data Figure 2) [54]. The sum of spikes in the *j*^*′*^*th* history window at time *t* in trial *k* is denoted by *h*_*j,k,t*_. The log of the instantaneous firing rate at time *t* and trial *k*, log *λ*_*k*_(*t*), is modeled as a linear combination of trial phase, *S*_*t*_ = *{s*_*t*,1_, *s*_*t*,2_, …*s*_*t,R*_*}*, and spike history *H*_*k,t*_ = *{h*_*t*,1_, *h*_*t*,2_, …*h*_*t,J*_ *}*,

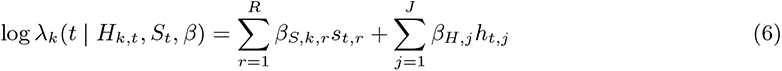

where the coefficients *β* = *{β*_*S*_, *β*_*H*_ *}* characterize the relationship between each covariate and the instantaneous firing rate. Alternately, we can express *λ*_*k*_(*t*) as

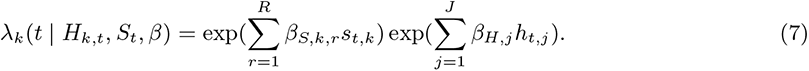

We assume that the inherent properties of the neuron remain constant across trials. Thus, the variation in firing rate across trials is modeled by a trial varying coefficient for the trial phase covariate, *β*_*S,k,r*_. We assume that the effect of trial phase modulation varies smoothly across trials, and so we link each *β*_*S,k,r*_ with a first order Gaussian process model:

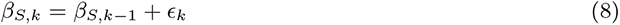

where *ϵ*_*k*_ is an *R*-dimensional Gaussian random variable with zero mean and covariance matrix Σ. The maximum likelihood estimates of 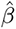 and 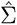 were computed with the EM algorithm [8, 46]. The median and 95% confidence intervals for quantities computed from 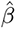, including 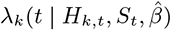 are computed via Monte Carlo [8].

Across a given trial, the terms 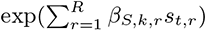 for *t* = [1, *T*] characterize within-trial firing rate variation (the task encoding effect), as well as the trial-average firing rate. To isolate task encoding effect, we compute 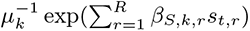, where *µ*_*k*_ is the trial average firing rate, 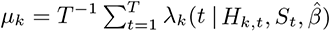. The task encoding effect enables comparison of how trial information is encoded between trials with drastically different trial-average firing rates. To compare the task encoding effect between experimental periods (baseline, inhibition, and recovery), we used computed the median and 95% confidence intervals for the mean task encoding effect across 20 trials within each period via Monte Carlo [8].

We estimated the model parameters from odd (training) trials, and performed goodness of fit assessment on even (test) trials as described in [8] (Extended Data Figure 5). To assess if there was a significant task encoding effect, we compare the full model to a reduced model. The reduced model assumes the base rate within trials is constant; that is, there is no variation firing rate due to the task. Thus, this model is a reduced version of the SS-GLM described above, where K trial covariates, *β*_*red,S*_ = *{β*_*red,S*,1_, *β*_*red,S*,2_, …, *β*_*red,S,K*_ *}*, capture rate variation across trials:

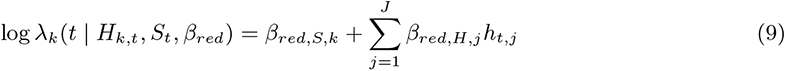

and the coefficients *β*_*red*_ = *{β*_*red,S*_, *β*_*red,H*_ *}* characterize the reduced model. As above, *β*_*red,S,k*_ varies across trials according to a Gaussian process model:

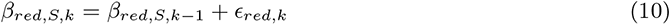

where *ϵ*_*red,k*_ is a 1-dimensional Gaussian random variable with zero mean and variance *σ*^2^. The significance of the task encoding effect was then assessed using the Likelihood Ratio Test, with the test statistic *λ*_*LR*_,

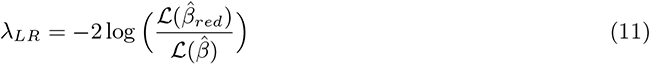

where 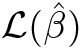 denotes the likelihood of the full model and 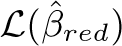 denotes the likelihood of the reduced model. Under the null hypothesis that *β ∈ β*_*red*_, *λ*_*LR*_ has been shown to follow a *χ*^2^ distribution with *ν* degrees of freedom, where *ν* is equal to the difference in the number parameters between the full and the reduced models. In this case, *ν* = (*K* + 1) × (*R −* 1). Thus, the probability of the null hypothesis that modeling within trial variation in firing rate does not provide a significantly improved characterization of single unit activity can be measured with *p* = 1 *− F* (*λ*_*LR*_|*ν*), where *F* (·|*ν*) denotes the cumulative distribution function of the *χ*^2^ distribution with *ν* degrees of freedom.

To investigate task encoding between different trial variants, we further separated the training trials based on which trial variant they belonged to (e.g., correct vs. incorrect trials), and estimated a unique set of model parameters for each trial variant. The identity of a given test trial was decoded by comparing the deviance of the observed test data from each estimated model. Deviance is a generalization of sum of squared error (used to assess the goodness-of-fit of Gaussian linear models) for generalized linear models. For example, to decode between correct and incorrect trials using model estimates 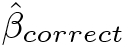 and 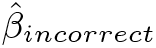 computed from correct and incorrect training trials, respectively, we considered spike train *n*_*k*_ = [*n*_*k*,1_, *n*_*k*,2_, …, *n*_*k,t*_] for trial *k*, where *t* denotes within trial time and *n*_*k,t*_ = 1 if there is a spike at time *t* in trial *k*. 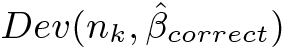 and 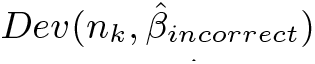 define the deviance of *n*_*k*_ from the correct and incorrect trial models, respectively. If 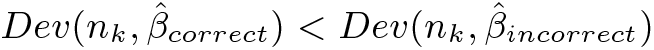, then we decoded the identity of trial *k* as correct. To measure how decoding accuracy changed throughout the experiment, we calculated the decoding accuracy across sliding windows of 50 trials, where decoding accuracy is the fraction of trials where the decoded trial variant matches the true trial variant. To estimate 95% confidence intervals of the decoding accuracy, we repeated the above approach across *I* = 1000 replicates of 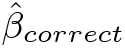 and 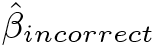. We compared decoding accuracy to a null decoder which assigns correct vs. incorrect trials according to the underlying behavioral accuracy. That is, if the probability of a trial being correct was *p*_*correct*_ = 0.76, any given test trial would be assigned as correct 76% of the time. The proportion of trials that would be accurately classified with the null decoder can be computed as follows:

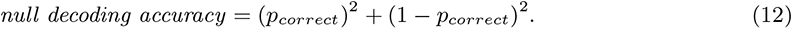

#### 4.3.3 Local Field Potential Analysis

##### 4.3.3.1 Time-frequency analysis of LFP Oscillations

To compare awake LFP oscillatory structure before and after GABA or saline delivery, we characterized the LFP spectra using the fitting oscillations and one over F (FOOOF) algorithm [42]. The FOOOF algorithm models a given power spectral density (PSD), as a sum of an aperiodic exponential decay component and gaussian peaks for each periodic oscillation,

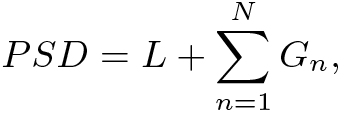

where *N* is the total number of periodic oscillations in the spectra, *G*_*n*_ is a gaussian peak centered at the frequency of the periodic oscillation, and *L* = *b −* log(*F* ^*χ*^) defines the aperiodic component. The aperiodic component has a broadband offset, *b*, and a rate of exponential decay, *χ*, which characterizes the exponential decay in power with increasing frequency, *F*. Each parameter is estimated in an iterative, unsupervised algorithm. We used the following hyperparameters as inputs to the algorithm: minimum peak width = 5 Hz, maximum peak width = 15 Hz, minimum peak height = 0.1, maximum number of peaks = 6.

We used bootstrapping to estimate the distribution of mean FOOOF parameter values for each experimental condition. For each experimental condition, (*M ×* 60 *× I*) 30 second LFP segments were randomly sampled from 10 minutes before and after microinfusions, where *M* is the number of sessions for the given experimental condition (*M* = 4 premotor cortex recordings with GABA infusion, *M* = 3 premotor cortex recordings with saline infusion, *M* = 1 putamen cortex recording with GABA infusion), and *I* = 1000 is the number of bootstrap samples used to estimate mean parameter values. We then calculated the multitaper spectra (time half-bandwidth product = 10, n tapers = 19) from each sample of LFP data. The multitaper spectra were used as inputs to the FOOOF algorithm. From FOOOF, we extracted periodic spectral peak frequencies and their corresponding power, the exponent and offset of the aperiodic component of the spectra, and the estimated spectra itself for each of the LFP samples. Each spectra had between 2–6 spectral peaks. To identify frequency ranges of dominant spectral peaks, we performed k-means clustering on peak frequencies of samples collected during the baseline recording periods for the premotor cortex and putamen. We defined each frequency range as the 2.5th and 97.5th percentiles of the peak frequencies assigned to the corresponding cluster, rounded to the nearest integer. From the premotor cortex recordings, we identified seven frequency ranges with periodic oscillations: 5–8 Hz, 11–17 Hz, 17–25 Hz, 27–35 Hz, 35–47 Hz, 47–62 Hz and 70–100 Hz. From the putamen recordings, we also identified seven frequency ranges: 5–8 Hz, 10–16 Hz, 17–22 Hz, 27–32 Hz, 34–42 Hz, 54–60 Hz, and 76–83 Hz. We defined the power of oscillations in each frequency range as the peak power of oscillations identified within that frequency range; the power was zero if no oscillations were detected in a given range (there was no power above the aperiodic component). We calculated the mean power in each frequency band and mean aperiodic parameters across (*M ×* 60) spectral samples from each experimental condition. We repeated this process across each of the *I* = 1000 bootstrap samples in order to generate 1000 estimates of the mean value of each FOOOF parameter. We then calculated the median and 95% confidence intervals for the difference in mean parameter value before and after microinfusions by extracting the 50^*th*^, 2.5^*th*^ and 97.5^*th*^ percentiles.

##### 4.3.3.2 Characterizing task-evoked LFP

We modeled the relationship between LFP amplitude and trial epoch using an autoregressive model with trial covariates,

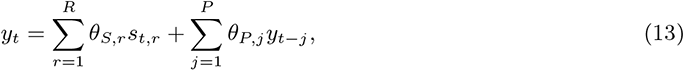

where *y*_*t*_ is the LFP amplitude at time *t*. As in the spike model, the trial subphase covariate, *S*_*t*_ = *{s*_*t*,1_, *s*_*t*,2_, …*s*_*t,R*_*}* is a set of *R* unit pulse functions, where *s*_*r,t*_ = 1 if time *t* is within trial subphase *r. P* defines the model order of the AR process. Here, we use *P* = 2, which is sufficient to estimate the basic structure of the LFP spectrum [55]. The coefficients *θ* = *{θ*_*S*_, *θ*_*P*_ *}* characterize the AR model where LFP amplitude varies with task phase.

As with the spike model, we estimated the LFP model from odd (training) trials and performed goodness of fit assessment on even (test) trials. Goodness of fit was assessed by computing the autocorrelation function of the residuals, which should be white noise if the model is an accurate description of the data. In contrast to the spike model, we found that models with stationary trial subphase coefficients had white noise residual autocorrelation, which reflects a stationary relationship between the trials and LFP amplitude (Extended Data Figure 10). To decode between trial variants from LFP, we employed a similar method to the approach used for spike data. First, we estimated models from training trials belonging to each trial variant. For example to decode between correct and incorrect trials, we estimated 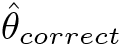 and 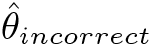 from correct and incorrect training trials, respectively. To account for changes in the LFP spectra (Figure 3, Figure 5h,i), we separately estimated 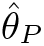 for each test trial. Then, for a given test trial, we calculated the sum of squared error (SSE) between observed and predicted LFP from models trained with each trial variant, and selected the trial variant with the lowest SSE. (In the case of a linear gaussian model, deviance generalizes to SSE.) To estimate 95% confidence intervals of the decoding accuracy, we repeated the above approach across *I* = 1000 replicates of 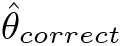 and 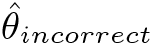. We defined the null decoding accuracy as described in Section 4.3.2.

#### 4.3.4 Behavior Analysis

Behavioral task accuracy was defined as the number of correct trials divided by the total number of complete trials; trials where the macaque did not fixate throughout the fixation, sample, and delay phases were excluded from analysis. To determine if there was a change in task accuracy before and after microinfusions, we considered two comparisons: (1) the task accuracy after a microinfusion compared to the baseline accuracy for the same session, and (2) task accuracy after a microinfusion compared to the baseline accuracy across all sessions. We estimated median and 95% confidence intervals for the change in task accuracy by generating *I* = 10000 bootstrap samples from each set of trials, measuring the difference in task accuracy between post-GABA and pre-GABA bootstrap samples, and extracting the 50^*th*^, 2.5^*th*^ and 97.5^*th*^ percentiles of the differences. We considered a change in task accuracy to be significant only if the 95% confidence intervals for the change in task accuracy did not include zero for both within session (1) and across session (2) baseline comparisons (Extended Data Figure 8).

## 5 Acknowledgments

The authors thank the MIT Division of Comparative Medicine for their expert support of this work. This work was funded in part by the National Institutes of Health (ICG: 5T32EB019940-05; ENB, EKM: P01- GM118629), the National Science Foundation (ICG: GRFP-1745302; PA: DMR-1419807, EEC-1028725), the National Institute of Mental Health (EKM: R37MH087027), the Office of Naval Research (EKM: MURI N00014-16-1-2832), the McGovern Institute for Brain Research (PA), and the Picower Institute for Learning and Memory (ENB, EKM). ICG is a recipient of the MIT SoE MathWorks Fellowship, and AJM is a recipient of the Simons Center for the Social Brain Postdoctoral Fellowship.

## Declarations

ENB has licensed IP to Masimo for the analysis of EEG under anesthesia. ENB is a cofounder of PASCALL, a company constructing control systems for anesthesia.

