## Extended Data for "Multifunctional fibers enable modulation of cortical and deep brain activity during cognitive behavior in macaques"

### 6 Extended Data

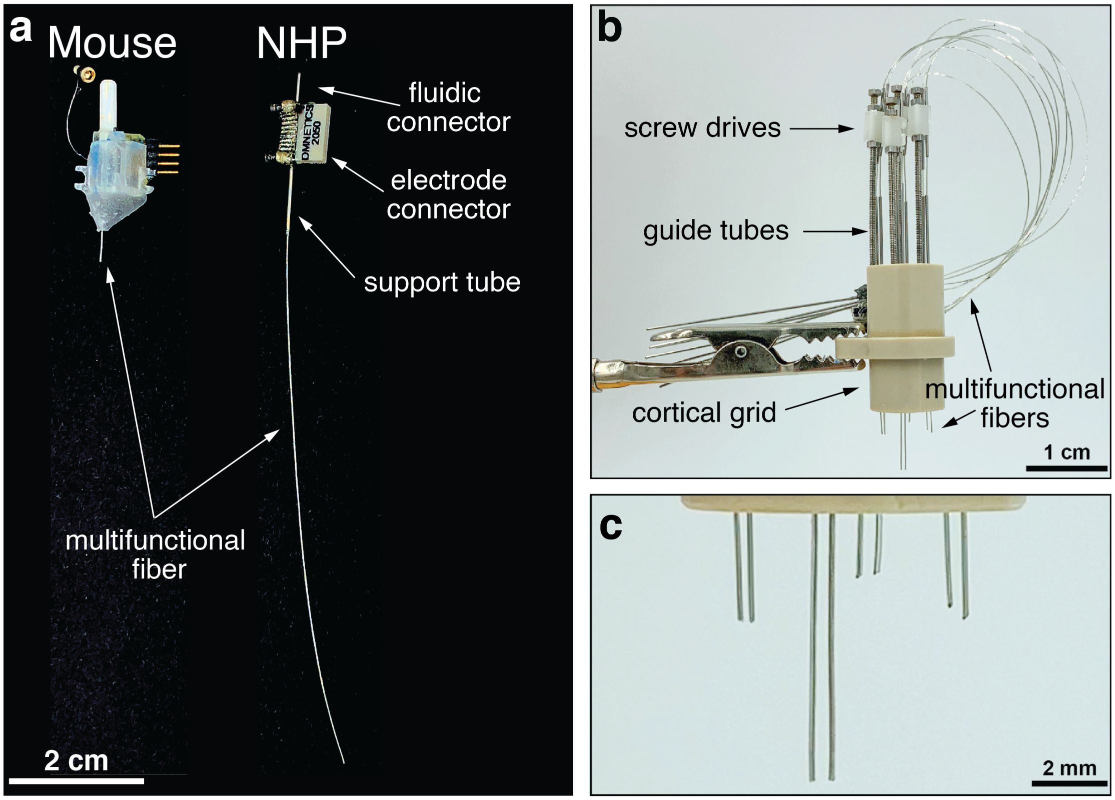


Extended Data Figure 1: **Multifunctional fiber-based neural interfaces.** **a,** Side-by-side comparison of the form factor of multifunctional fibers used for mouse vs. NHP research. **b,** An array of multifunctional fibers arranged in a cortical grid with four independently actuated screw drives. The fibers are fixed to the white shuttles at the top of the screw drive, and the fibers extend to a backend connector fixed to the side of the grid. Fixing the fibers to the shuttle enables precise depth control, while the flexibility of the fibers allows for unique backend configurations. **d,** Independently actuated screw drives enable implantation of multiple fibers at varying depth across a 1×1 cm$^2$ area.

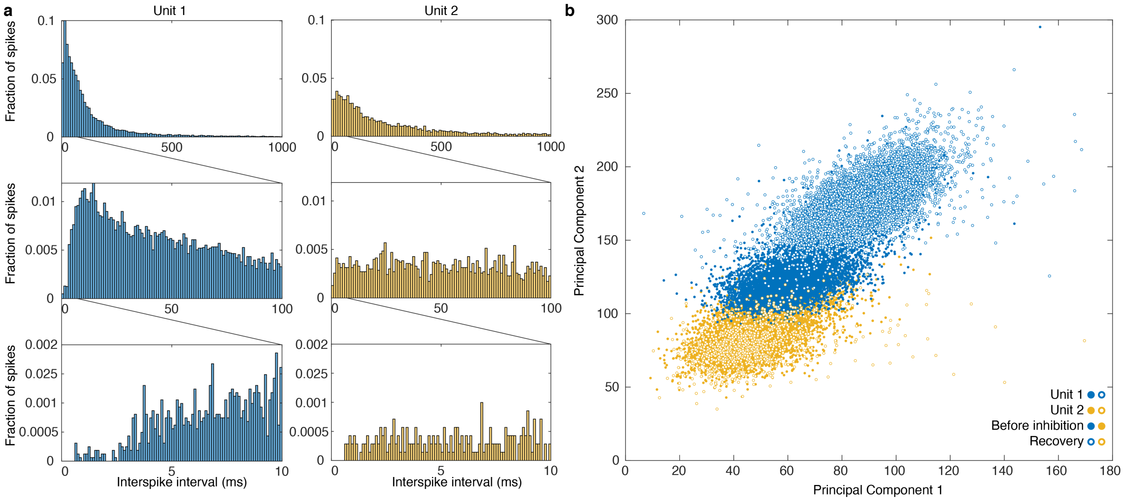


Extended Data Figure 2: **Properties of units in Figure 2.** **a,** Histograms of the interspike intervals of unit 1 and unit 2, where the y-axis is the proportion of spikes in each bin. **b,** Principle components of unit 1 (L-ratio = 0.13) and unit 2 (L-ratio = 0.30) before and after GABA. The principle components of unit 1 were stable throughout the experiment, while there was a slight shift in unit 2. The shift is due to the increase in amplitude on channel 1 during the recovery period.

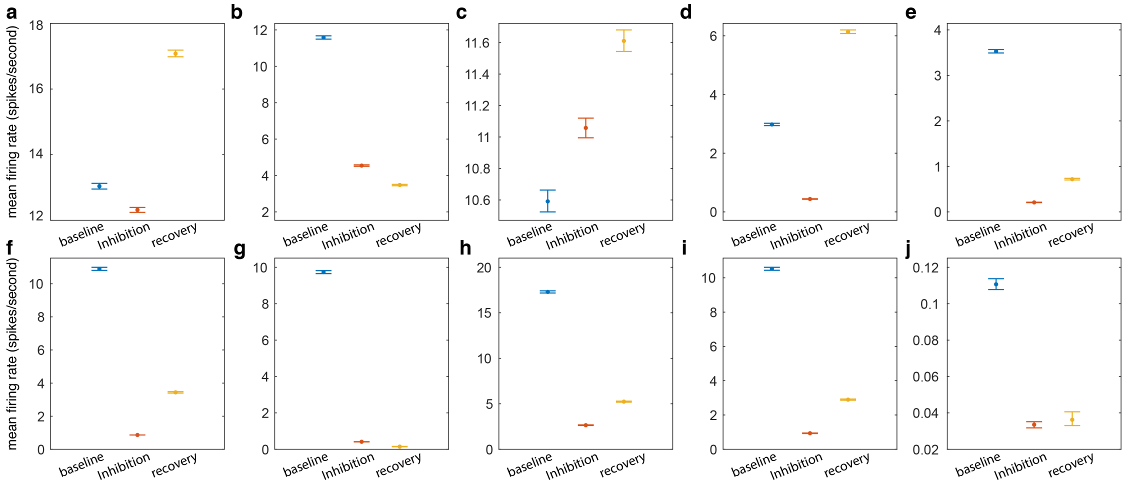


Extended Data Figure 3: **All units' responses to GABA microinfusions in the premotor cortex.** The mean firing rate of trials in each experimental period (baseline, inhibition, and recovery) and 95% confidence intervals are plotted across the 10 units recorded across 4 GABA sessions. The units corresponding to panel d and e are units 1 and 2 in Figure 2, respectively, and the activity of the unit corresponding to panel f is shown in Figure 4.

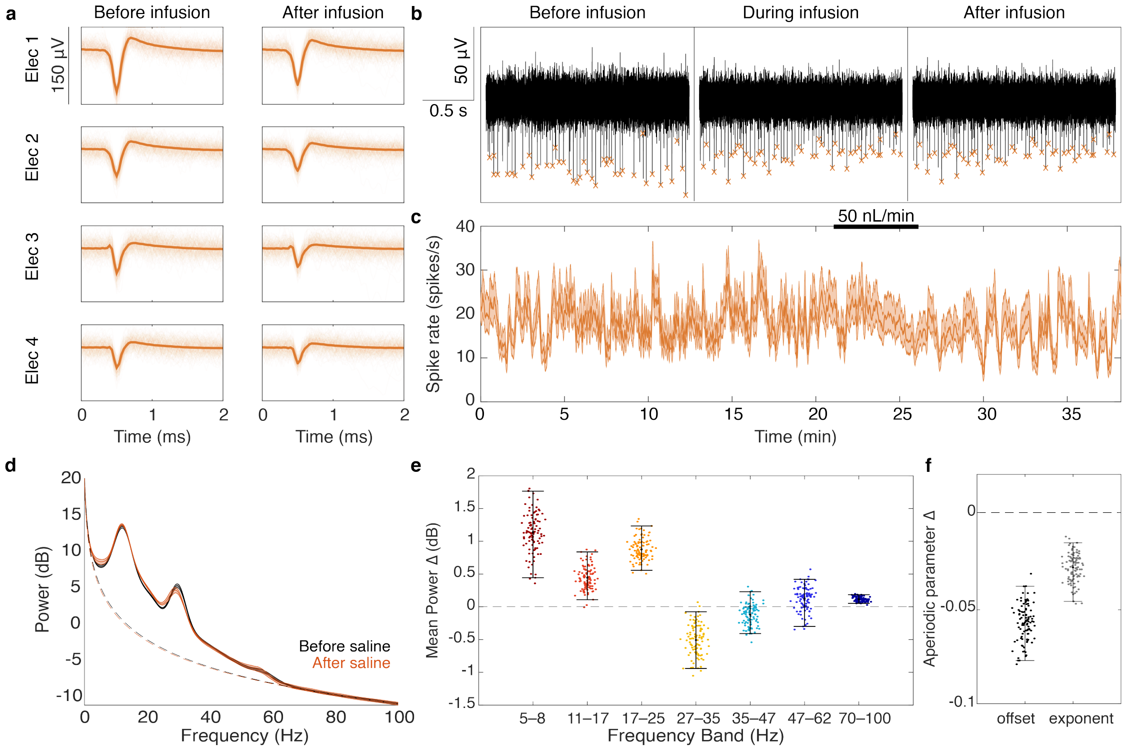


Extended Data Figure 4: **Measuring the effect of the saline vehicle on neurophysiology a**, The waveform of single unit activity was stable in both shape and relative amplitude before and after 50 nL/min infusions of saline. **b**, The noise floor of the recordings was not affected by intracortical saline delivery, allowing for consistent identification of single unit waveforms before, during, and after the 50 nL/min saline infusion. The orange x's mark spikes from the unit shown in panel a. **c**, The firing rate of single unit activity was modeled with a state-space point-process model (Methods). 95% confidence intervals for the estimated rate are shown with orange shading. There were no significant downstates following saline infusion. **d**, The mean and 95% confidence intervals of 100 bootstrap samples of the mean spectra calculated from 10-minute periods before and after saline are indicated in black and red, respectively, **e**, Scatter plots of 100 bootstrap estimates of the mean peak power in the 10 minutes before the saline infusion vs. the 10 minutes after the saline infusion. There was a significant increase in 4–8, 11–17, 17–25, and 70–100 Hz power, and a significant decrease in 27–35 Hz power. **f**, The mean estimated offset and exponent of the aperiodic component were decreased following saline infusion.

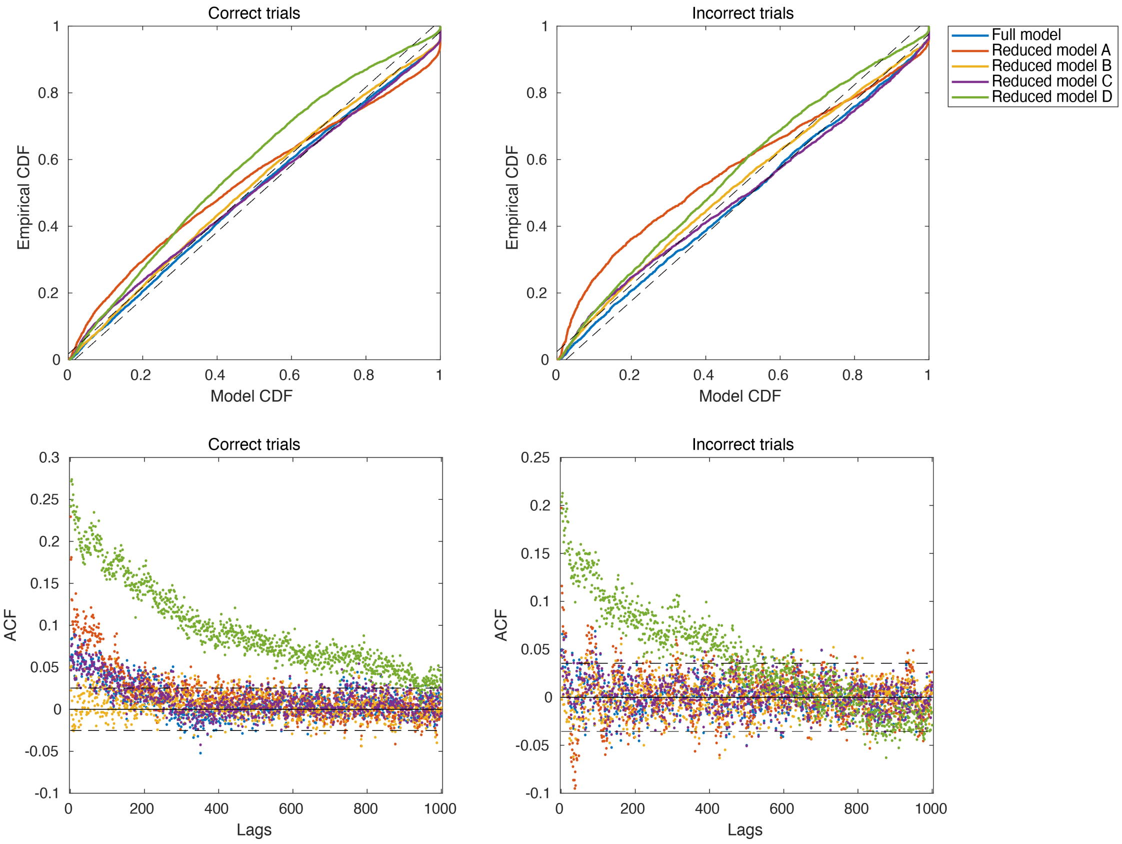


Extended Data Figure 5: **Goodness of fit of candidate GLM models for characterizing dynamic task-related activity of the unit presented in Figure 3.** Candidate models included the full model: SS-GLM with history coefficients and task phase coefficients that varied across trials, Reduced model A: SS-GLM with no history coefficients and a single task phase parameter that varied across trials, Reduced model B: SS-GLM with history covariates and a single task phase parameter that varied across trials, Reduced model C: SS-GLM with task phase coefficients that varied across trials and no history covariates, Reduced model D: stationary GLM (GLM with history coefficients and task phase coefficients that were constant across trials). Top row: KS-plots for rescaled interspike intervals for correct and incorrect trials. In the case of an exact model fit, the rescaled interspike intervals would follow the 45° line. The dashed black lines define 95% confidence bounds of an exact model fit. Bottom row: the autocorrelation function (ACF) of the rescaled interspike intervals for correct and incorrect trials. In an exact model fit, the rescaled interspike intervals would be independent and ACF = 0 for all lags. The dashed black lines define 95% confidence bounds of an exact model fit.

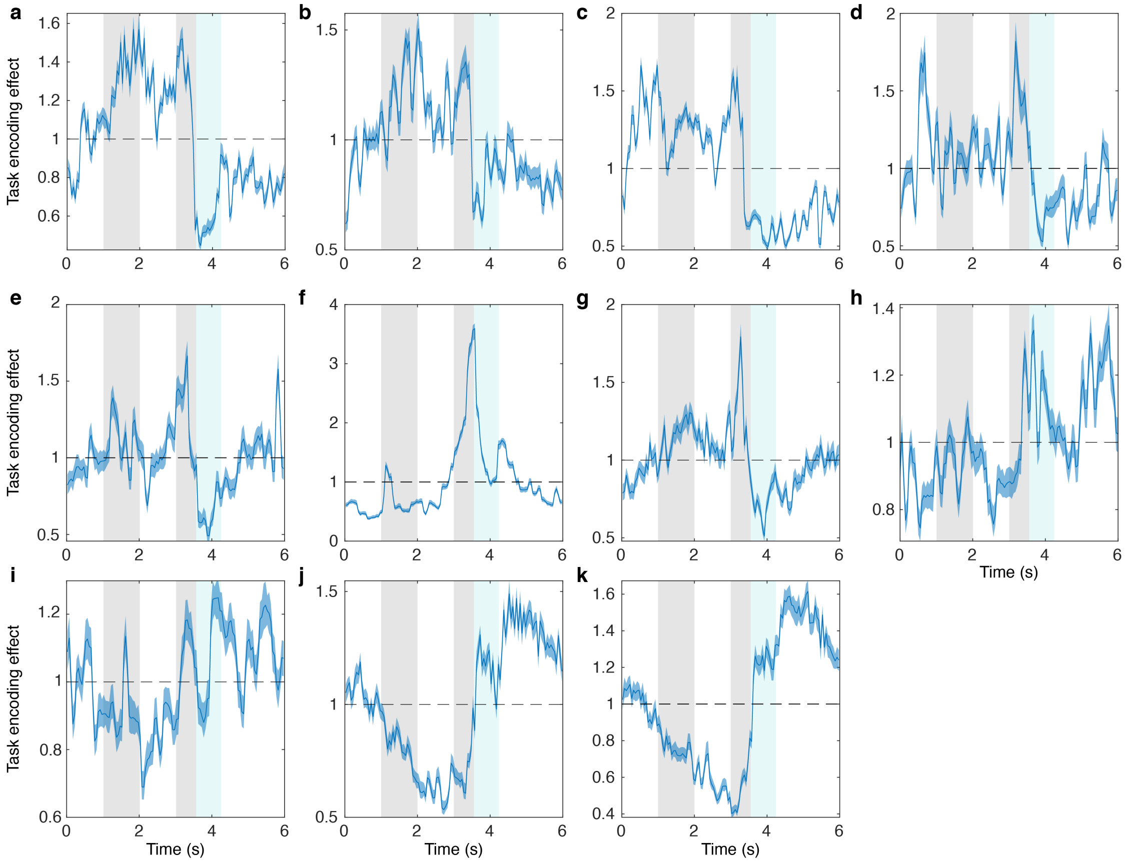


Extended Data Figure 6: **All premotor cortex units' evoked activity.** Over 4 GABA sessions and 3 saline sessions, there were 11 units whose firing rate varied significantly within trials (Likelihood Ratio test, $p < 0.05$, Methods). The first row shows units that had elevated firing rates durin the fixation-match phases of the trial, followed by a sharp decrease in firing rate during the reward period. The second row shows units whose firing rate peaked during the match phase. Row three shows units with elevated firing rate during and following the reward phase. The units corresponding to panel d and e are unit 1 and 2 in Figure 2, respectively, and the unit corresponding to panel f is shown in Figure 4.

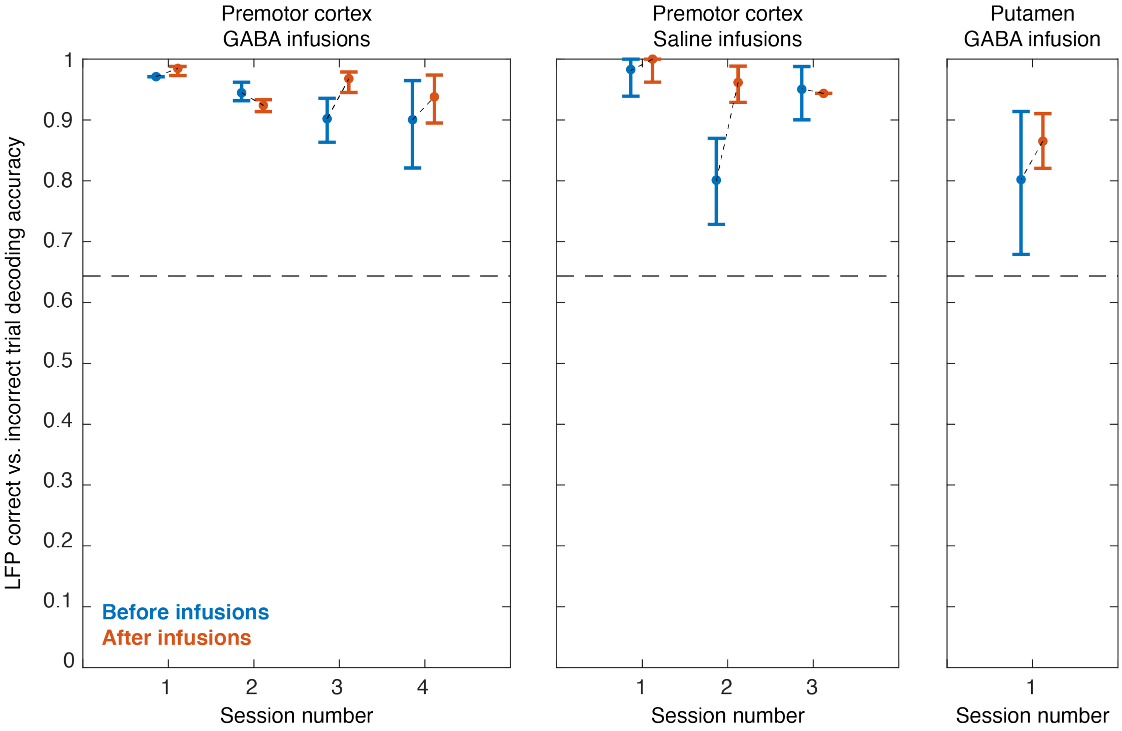


Extended Data Figure 7: **Change in LFP decoding accuracy due to microinfusions.** The change in LFP decoding accuracy before and after microinfusion was measured from 4 premotor cortex GABA sessions, 3 premotor cortex saline sessions, and 1 putamen GABA session. The null decoding accuracy across all sessions, 64%, is indicated with the horizontal dotted line.

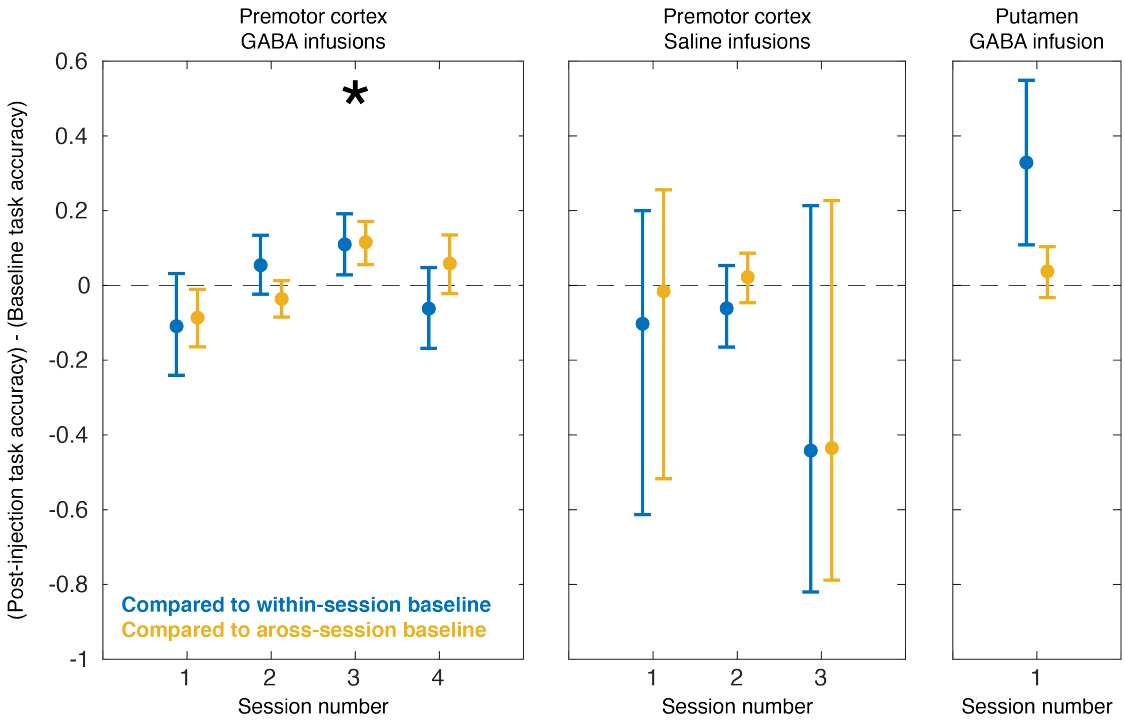


Extended Data Figure 8: **Behavioral change due to microinfusions.** The change in task accuracy before and after microinfusions was measured from 4 premotor cortex GABA sessions, 3 premotor cortex saline sessions, and 1 putamen GABA session. Task accuracy after microinfusions was compared to task accuracy at baseline (before GABA) from the same session and task accuracy across the baseline periods of all sessions. There was 1 premotor GABA session with significantly increased task accuracy for both comparisons. There was an increase in task accuracy compared to the within-session baseline for the putamen session, but not compared to the across-session baseline. Thus, the increase in task accuracy can be explained by lower than normal task accuracy during the putamen baseline period.

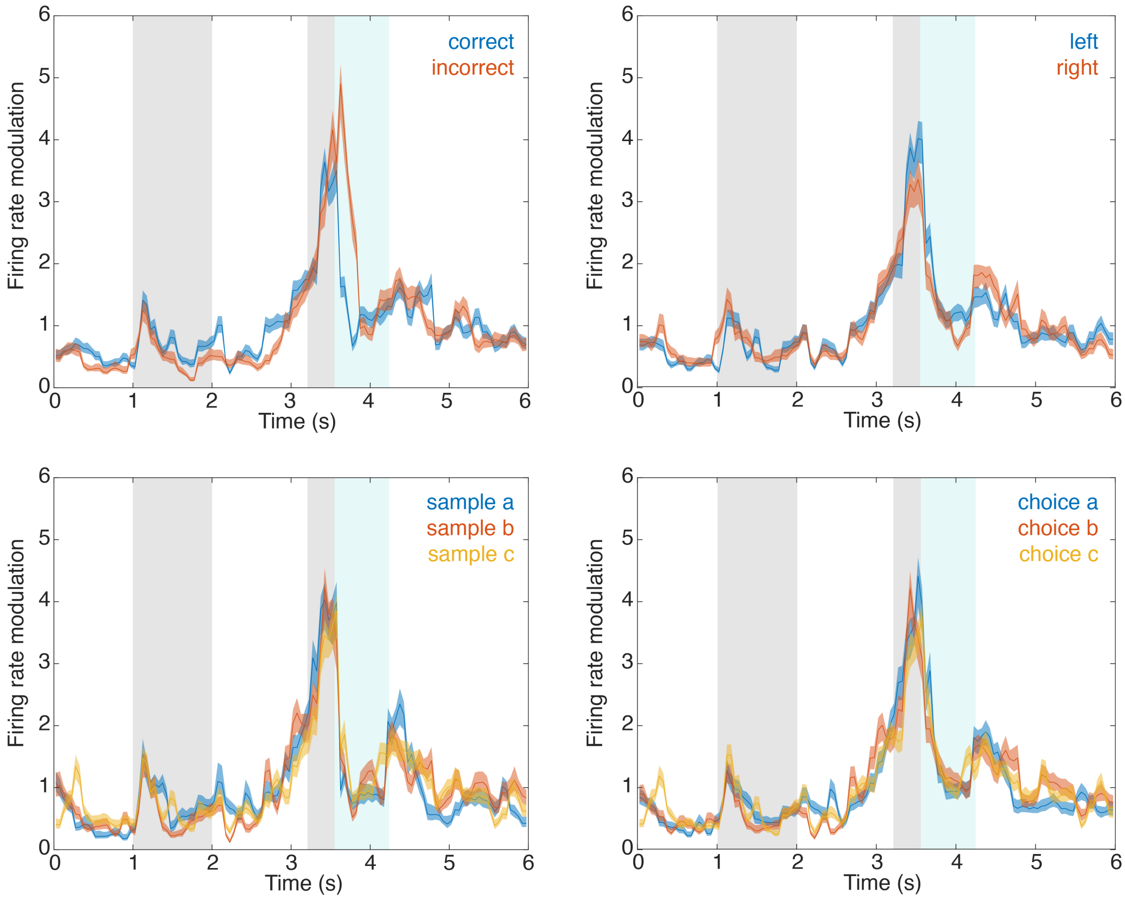


Extended Data Figure 9: **Comparing encoding across trial variants.** We estimated variant specific SS-GLM models by training 2 or 3 models with trials from each variant. For instance, to determine if there was a difference in task encoding between correct and incorrect trials, we trained one model with odd trials that were correct, and another with odd trials that were incorrect. We then compared the task encoding effect from 20 baseline trials to determine if there was a significant difference in task encoding between trial variants. For the unit shown in Figure 4, we investigated **a,** correct vs. incorrect trial encoding, **b,** left vs. right saccade encoding, **c,** sample ID encoding, and **d,** choice ID encoding. We found that there was the largest between-variant difference in correct vs. incorrect trial encoding.

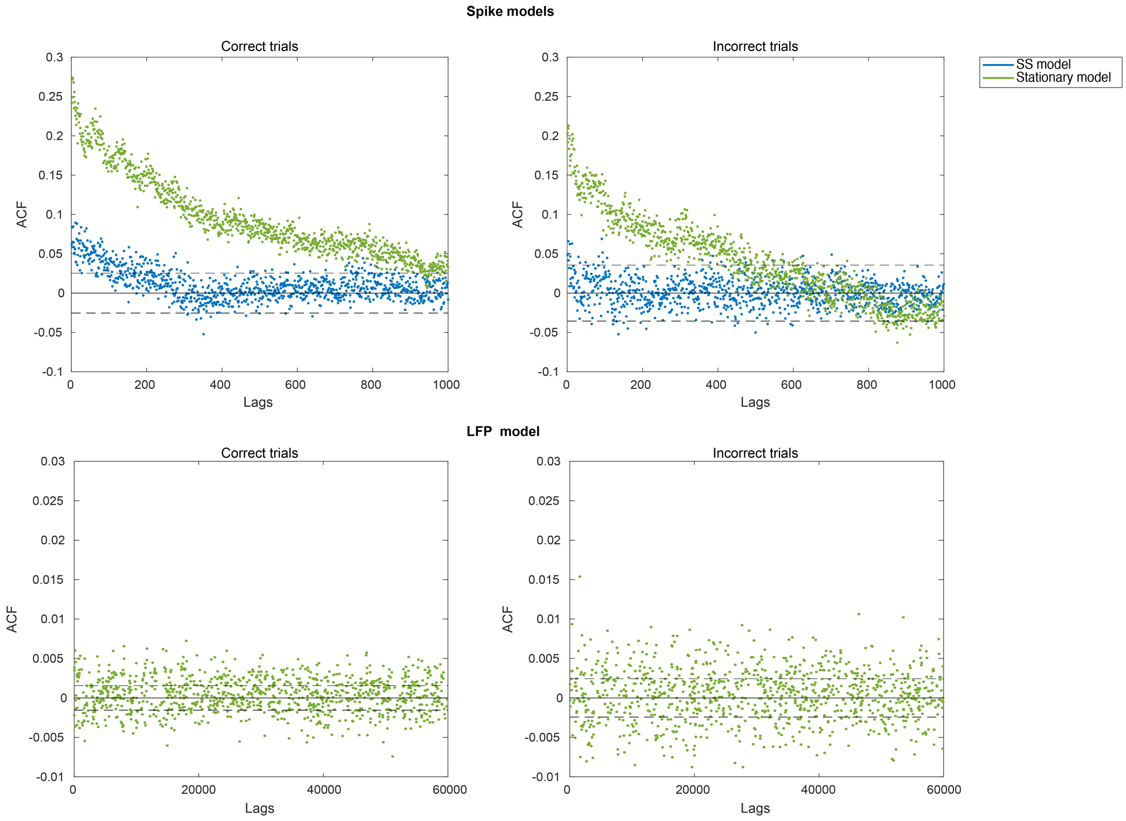


Extended Data Figure 10: **Autocorrelation functions (ACF) for state-space vs. stationary models for spike and LFP evoked activity** Top row: The ACF for rescaled interspike intervals for correct and incorrect trials computed from the spike models. The blue dots indicate rescaled interspike intervals estimated from the full SS-GLM model (SS-GLM with history coefficients and task phase coefficients that varied across trials) and green dots indicate rescaled interspike intervals estimated from the stationary GLM model (GLM with history coefficients and task phase coefficients that were constant across trials). In an exact model fit, the rescaled interspike intervals would be independent and ACF $= 0$ for all lags. The dashed black lines define 95% confidence bounds of an exact model fit. Note that there is significant structure to the ACF of rescaled interspike intervals estimated from the stationary model, which is mitigated by addressing cross-trial variability in firing rate with the SS-GLM model. Bottom row: The autocorrelation function of residuals for correct and incorrect trials computed from the LFP AR(2) model with task phase coefficients. Note that there is not significant structure to the ACF, indicating that a stationary model is sufficient to characterize LFP across trials. The maximum number of lags was chosen to capture temporal relationships across up to 10 trials.
